# DNA methylation dynamics during stress-response in woodland strawberry (*Fragaria vesca*)

**DOI:** 10.1101/2022.03.04.483002

**Authors:** María-Estefanía López, David Roquis, Claude Becker, Béatrice Denoyes, Etienne Bucher

**Affiliations:** Crop Genome Dynamics Group, Agroscope, 1260 Nyon, Switzerland; Department of Botany and Plant Biology, Faculty of Sciences, University of Geneva, Geneva, Switzerland; LMU BioCenter, Faculty of Biology, Ludwig-Maximilians-University Munich, D-82152 Martinsried, Germany; Univ. Bordeaux, INRAE, Biologie du Fruit et Pathologie, F-33140 Villenave d’Ornon, France

**Keywords:** epigenetics, stress response, transposable elements, transcription factors, centromeres

## Abstract

- Environmental stresses can result in a wide range of physiological and molecular responses in plants. These responses can also impact epigenetic information in genomes especially at the level of DNA methylation. DNA methylation is the hallmark heritable epigenetic modification and plays a key role in silencing transposable elements (TEs). Although DNA methylation is an essential epigenetic mechanism, fundamental aspects of its contribution to stress responses and adaptation remain obscure.
- We investigated epigenome dynamics of wild strawberry (*Fragaria vesca*) in response to variable environmental conditions at DNA methylation level. *F. vesca* methylome responded with great plasticity to ecologically relevant abiotic and hormonal stresses. Thermal stress resulted in substantial genome-wide loss of DNA methylation. Notably, all tested stress conditions resulted in marked hot spots of differential DNA methylation near centromeric or pericentromeric regions, particularly in non-symmetrical DNA methylation context. Additionally, we identified differentially methylated regions (DMRs) within promoter regions of transcription factor (TF) superfamilies involved in plant stress-response and assessed the effects of these changes on gene expression.
- These findings improve our understanding on stress-response at the epigenome level by highlighting the correlation between DNA methylation, TEs and gene expression regulation in plants subjected to a broad range of environmental stresses.

## Introduction

Plants in natural environments are exposed to multiple stimuli, including numerous biotic and abiotic stresses that make it necessary for plants to develop strategies to rapidly adapt. According to the Global Climate Report 2020, the past 10 years were the warmest recorded around the globe in our era. The greater temperature variability has resulted in both droughts and extreme precipitations, affecting not only natural plant populations but also crop production (WMO, 2021). In order to face these challenges, we need to better understand the mechanisms which allow plants to rapidly adapt and evolve to better cope with increasing climate change-related stresses. Recent advances in genome sequencing have revealed how dynamic plant genomes can be under stressful scenarios (Kersey, 2019; Nguyen *et al.*, 2019, Roquis 2021). This dynamism can be attributed to both genetic and epigenetic mechanisms which can contribute to specific traits (Varotto *et al.*, 2020; Wang *et al.*, 2019). However, how epigenetic information is influenced by stresses (Quadrana & Colot, 2016; Lämke & Bäurle, 2017; MacKelprang & Lemaux, 2020; Varotto *et al.*, 2020) and whether these can contribute to adaptation requires a better understanding. DNA methylation is an epigenetic mark which exists in three sequence contexts in plants: CG, CHG, and CHH (H = A, C, or T). Each of them is regulated by distinct, but also interconnected silencing mechanisms (Law & Jacobsen, 2010; Sahu *et al*., 2013; Matzke & Mosher, 2014). Symmetric methylation in the CG sequence context (mCG) has been found to be enriched in gene bodies but the biological function of gene body methylation (gbM) remains unclear (Bewick & Schmitz, 2017; Bewick *et al.*, 2019). mCG is highly heritable and able to persist over many generations. Conversely, DNA methylation in the CHG (mCHG) and CHH (mCHH) sequence contexts show a lower stability (Becker *et al.*, 2011; Kuhlmann *et al.*, 2014; Schmitz *et al.*, 2011; Williams & Gehring, 2017). DNA methylation is a hallmark epigenetic modification, contributing to the regulation of many biological processes such as genome stability, definition of euchromatin and heterochromatin, control of gene expression, and, most importantly, silencing of transposable elements (TEs) (Bewick & Schmitz, 2017; Bucher *et al.*, 2012; Zhang *et al*., 2018). Studies of DNA methylation variability in natural *Arabidopsis* accessions have shown a clear correlation between epigenomic changes in coding and non-coding genomic regions and environmental stimuli, suggesting a role for DNA methylation in adaptation (Kawakatsu *et al.*, 2016). More generally, it has been found that DNA methylation changes may be implicated in morphological changes in response to different climates in plants (Guarino *et al.*, 2015; González *et al.*, 2018). However, whether stress-induced methylome alterations at individual loci or across the entire genome contribute to heritable changes in DNA methylation patterns and adaptation remains uncertain.

A large fraction of the genome-wide observations on the functional properties of DNA methylation in unfavorable growth conditions have been carried out on Arabidopsis (van Dijk *et al.*, 2010; Colaneri & Jones, 2013; Jiang *et al.*, 2014; Shen *et al.*, 2014). However, its genome composition differs significantly from cultivated crops that are characterized by larger genomes such as wheat, maize or sugarcane (Niederhuth *et al.*, 2016; Vidalis *et al.*, 2016). Indeed, genomic DNA methylation content is very low in Arabidopsis (Alonso *et al.*, 2015). Woodland strawberry (*Fragaria vesca,* diploid, 219 Mb, 2n= 2x=14) (Edger *et al.*, 2018) is an interesting model for the study of DNA methylation because it regulates key developmental traits of this species, including seed dormancy (Zhang *et al.*, 2012) and fruit ripening (Cheng *et al*., 2018). A notable example concerning environmental impacts on DNA methylation is the documented loss of methylation at TEs at high altitudes and how it can contribute to local adaptation to local conditions in natural populations of wild strawberry (De Kort *et al*., 2020; Sammarco *et al.*, 2022).

Here, we wanted to assess how stresses influence DNA methylation in *F. vesca* which has a genome roughly twice as large as that of Arabidopsis, and at the same time offers advantages for functional genomic and epigenomic studies compared to genetically complex cultivated octoploid strawberry (*F. x ananassa*) (2n=8x=56) (Edger *et al.*, 2019; Shulaev *et al.*, 2011; Vitte *et al.*, 2014). We present high-resolution data describing the impact of a diverse set of stresses on DNA methylation in *F. vesca*. Depending on the stress conditions, we found that *F. vesca* can respond with global and/or local changes in DNA methylation. Notably, these changes impact key transcription factors (TFs) such us APETALA2/ethylene-responsive element binding factors (*AP2/EREBP*) but also stress specific TFs such as heat shock transcription factors (*HSF*s). Surprisingly, we find that all stresses have an impact on DNA methylation in centromeric or pericentromeric regions implying that these genomic regions may act as stress-responsive rheostats. Finally, we describe the DNA methylation dynamics at TE flanking regions as a strategy to maintain homeostasis during stress responses.

## Materials and Methods

### Plant growth and material

All strawberry plants used in this study were a homozygous cultivated near-isogenic line (NIL), Fb2:39–47, *F. vesca cv. Reine des Vallées* (RV), possessing the “r” locus on chromosome 2 which causes this accession to propagate vegetatively through stolon development (Urrutia *et al.*, 2015). Seeds from a single founder plant were germinated in water over Whatman filter paper for two weeks and transferred to 50% MS medium (Murashige & Skoog, 1962) (Duschefa cat# M0222), 30% sucrose, and 2% phytagel (Sigma-Aldrich cat# P8169) and grown for 4 weeks prior to stress.

### Stress assays

One-month-old seedlings on agarose plates were exposed to different stresses under long-day conditions (16 h light 24°C/8 h dark 21 °C) in plant growth chambers (Panasonic, phcbi: MLR-352/MLR-352H). The seedling age was optimized to assure stress tolerance. For salt and drought stress, one-month-old plantlets were transferred to MS media supplemented with 100 mM sodium chloride (NaCl) (Sigma-Aldrich, cat# S9888) and 5% polyethylene glycol (PEG-6000) (−0.05 MPa) (Sigma-Aldrich, cat# P7181), respectively. For cold and heat stresses, plants were initially grown as described above. The plates were then transferred to either 6°C or 37°C chambers. High light was induced by 20,000 lx of illuminance (460 *μ*mol s^-1^ m^-2^) and low light with 80% sunblock black net leading to 4,000 lx of illuminance (92 *μ*mol s^-1^ m^-2^). To simulate a hormone stress, MS medium was supplemented with 0.5 mM salicylic acid (SA) (Sigma-Aldrich, cat# 247588). All stress assays were carried out for 2 weeks with 2 recovery days after one week. For sampling, roots were removed from 5 pooled plants for each treatment group (3 biological replicates) and harvested in 1.5 mL tubes between 9:00 a.m. and 11:00 a.m. and immediately flash-frozen in liquid nitrogen and stored at −80°C.

### Genome sequencing and assembly NIL Fb2

Genomic DNA from strawberry plants was extracted by a Hexadecyltrimethylammonium bromide (Cetrimonium bromide, CTAB) modified protocol (Healey *et al.*, 2014) and purified with Agencourt AMPure XP beads (cat# A63880). Long-read sequencing was performed for genome assembly; Genomic DNA by Ligation (Oxford Nanopore, cat# SQK-LSK109) library was prepared as described by the manufacturer and sequenced on a MinION for 72 h (Oxford Nanopore).

### Reference genome polishing

Reads obtained from nanopore were filtered with Filtlong v0.2.1 (https://github.com/rrwick/Filtlong) using --min_mean_q 80 and --min_length 200. Cleaned reads were then aligned to the most recent version of the *F. vesca* genome v4.0.a1 (Edger *et al*., 2018), with the annotation of *F. vesca* genome v4.0.a2 downloaded from the Genome Database for Rosaceae (GDR) (https://www.rosaceae.org/species/fragaria_vesca/genome_v4.0.a2) (Jung *et al.*, 2019), using minimap2 v2.21 (Li, 2018) with parameters -aLx map-ont --MD -Y. The generated BAM file was then sorted and indexed with samtools v1.11 (Li *et al.*, 2009). We used mosdepth v0.3.1 (Pedersen & Quinlan, 2018) to verify that coverage on chromosomic scaffolds was over 50 X. Sniffles v1.0.12a (Sedlazeck *et al.*, 2018) with parameters -s 10 -r 1000 -q 20 --genotype -l 30 -d 1000 was used to detect structural variations larger than 30 bp. We observed that larger structural variants (SV) were most likely falsely identified due to misalignments in regions with gaps or Ns, therefore the VCF files obtained from Sniffles were sorted and filtered with BCFtools v1.14 (Danecek *et al*., 2021) to keep only SV with less than 200 kb, supported by 10 or more reads and with allelic frequencies above 0.8 to isolate homozygous changes. The complete filtering command used was “bcftools view -q 0.8 -Oz -i ‘(SVTYPE = “DUP” II SVTYPE = “INS” II SVTYPE = “DEL” II SVTYPE = “TRA” II SVTYPE = “INV” II SVTYPE = “INVDUP”) && %FILTER = “PASS” && FMT/DV>9 && SVLEN>29 && SVLEN<200000’”

From the VCF listing all the structural variants that we detected in our *F. vesca* accession, we generated a substituted genome version based on the reference *F. vesca* genome v.4.0.a2. The reference genome was first indexed with samtools faidx v1.11 (Danecek *et al*., 2021) and a sequence dictionary was generated with Picard CreateSequenceDictionary v2.25.6 (https://broadinstitute.github.io/picard). The VCF containing the SV produced from our Nanopore sequencing was also indexed with gatk (Van der Auwera GA & O’Connor BD, 2020) IndexFeatureFile v4.2.0.0 (https://gatk.broadinstitute.org/hc/en-us/articles/360037262651-IndexFeatureFile). FastaAlternateReferenceMaker v4.2.0.0 (https://gatk.broadinstitute.org/hc/en-us/articles/360037594571-FastaAlternateReferenceMaker) was then run with the reference genome and the VCF file to generate a substituted genome representative of our *Fragaria* accession (Fb2).

As substituting our genome with the detected structural variants changes genomic coordinates, we also corrected the public GFF genome annotation of *F. vesca* (Y, Pi, Gao, Liu, & Kang, 2019) using liftoff v1.6.1 (Shumate & Salzberg, 2021). Liftoff also detects and annotates duplications within the substituted genome.

Transposable elements annotation was carried out using the EDTA transposable element annotation pipeline v. 1.9.6 (Ou *et al.*, 2019) on the substituted genome using default parameters. The DOI for the *Fragaria vesca* Fb2:39–47 genome is: 10.5281/zenodo.6141713.

### Whole-genome bisulfite sequencing (WGBS)

A modified CTAB DNA extraction protocol was performed using frozen above-ground tissues (Healey *et al.*, 2014). DNA libraries were generated using the NEBNext Ultra II DNA Library Prep Kit (New England Biolabs, cat# E7103S) according to the manufacturer’s instructions with the following modification for bisulfite treatment. DNA was sheared to 350 bp using a Covaris S2 instrument. The bisulfite treatment step using the EZ DNA Methylation-Gold kit (Zymo Research, cat# D5007) was inserted after the adaptor ligation. After clean-up of the bisulfite conversion reaction, library enrichment was done using Kapa Hifi Uracil+ DNA polymerase (Kapa Biosystems, cat# KK1512) for 12 PCR cycles, using the 96 single-index NEBNext Multiplex Oligos for Illumina (New England Biolabs, cat# E7335S). Paired-end reads were obtained on an Illumina (150 bps) NovaSeq6000 instrument at Novogene (Hongkong, China).

### Processing and alignment of bisulfite-converted reads

Sequencing data was analyzed by data collection software from read alignment to DNA methylation analysis: Epidiverse/wgbs pipeline (Nunn *et al.*, 2021). The pipeline included quality control using FastQC v.0.11.9 (http://www.bioinformatics.babraham.ac.uk/projects/fastqc/) and Cutadapt v.3.5 (https://github.com/marcelm/cutadapt/). Genome mapping was performed using erne-bs5 v.2.1.1 (<u>Prezza *et al.*, 2012</u>) with default parameters to generate the BAM files. Methylation calling and methylation bias correction was performed with Methyldackel v.0.6.1 (https://github.com/dpryan79/MethylDackel) with only uniquely-mapping reads. The pipeline used Nextflow v20.07.1 to run multitask in parallel. Because plant chloroplast DNA are not methylated (Fojtová *et al.*, 2001), reads originating from those sequenced were used to evaluate the bisulfite conversion rate. The pipeline is available at https://github.com/EpiDiverse/wgbs. An average of 82,771,701 reads (~ 50X coverage) were produced per sample, of which 81% mapped properly to the *F. vesca* genome. The average non-bisulfite conversion rate among the samples was 0.10 (See Table S1 for more details). To calculate global methylation ratios, output files from wgbs pipeline were pre-filtered for a minimum coverage of 5 reads using awk command and only the common cytosine positions were kept among all the samples using bedtools 2.28.0. The data were tested for statistical significance with an unpaired Student’s t-test. p < 0.05 was selected as the point of minimal statistical significance in all the analyses. R-packages ggplot2 v.3.3.5 and gplots v.3.1.1 were used for the visualization of the results.

The Bisulfite-sequencing data from this study have been submitted to European Nucleotide Archive (ENA, www.ebi.ac.uk/ena/, accessed on ERP135585) under the project PRJEB50996, raw read fastq accessions under ERR8684931:ERR8684954.

### Identification of differentially methylated regions (DMRs)

First, bedGraph files from wgbs pipeline were pre-filtered for a minimum coverage of 5 reads using awk command. These output files were then used as input for the EpiDiverse/dmr bioinformatics analysis pipeline for non-model plant species to define DMRs (Nunn *et al*., 2021) with default parameters (minimum coverage threshold 5; maximum q-value 0.05; minimum differential methylation level 10%; 10 as minimum number of Cs; Minimum distance (bp) between Cs that are not to be considered as part of the same DMR is 146 bp). The pipeline uses metilene v.0.2.6.1 (https://www.bioinf.uni-leipzig.de/Software/metilene/) for pairwise comparison between groups and R-packages ggplot2 v.3.3.5 and gplots v.3.1.1, for visualization results (Fig. S1). Based on our *F. vesca* genome transcript annotation and methylation data (overlapped regions with DNA methylation cytosines and DMRs), we detected the methylated genes, promoters, 3’ UTRs, 5’UTR and transposable elements in strawberry. Global DNA methylation and DMR plots were performed with R-package ggplot2. Gene analyses by methylation patterns and analysis of per-family TE DNA methylation profiles were performed with deepTools v.3.5.0 (Ramírez *et al.*, 2014). DMRs comparison between treatments were done by the Venn diagram v.1.7.0 R-package.

We produced several genome browsers tracks with DMRs that we integrated in our local instance of JBrowse available at the following url: https://jbrowse.agroscope.info/jbrowse/?data=fragaria_sub. The bed files of the DMRs can be downloaded here: 10.5281/zenodo.6141713

### Gene Ontology (GO) enrichment of differentially methylated genes

All methylated genes were annotated based on GO annotation downloaded from the Genome Database for Rosaceae (GDR) (https://www.rosaceae.org/species/fragaria_vesca/genome_v4.0.a2).

To better understand the potential function of the differentially methylated genes, GO functional classification of these genes was performed by AgriGO program v1.2 (Tian *et al.*, 2017). Genes and promoters were classified by genes contained hypo and hypermethylated DMRs. The GO slim library was used as reference GO reference type. Fisher’s exact test p-values were calculated for over-representation of the differential methylated genes in all GO categories and Hochberg (FDR) as multi-test adjust method. GO terms with p < 0.05 were considered as significantly enriched.

### Real-time Quantitative PCR Analysis

One-month-old seedlings after heat and salt stress assays (as described above) were collected. For sampling, roots were removed from 5 pooled plants for each treatment (3 biological replicates) and harvested in 1.5 mL tubes between 9:00 a.m. and 11:00 a.m. and immediately flash-frozen in liquid nitrogen and stored at −80°C. RNA was extracted using NucleoSpin RNA Plus, Mini kit for RNA purification with DNA removal column (Macherey-Nagel, cat# 740984.50). cDNA synthesis was performed using EvoScript Universal cDNA Master (Roche, cat#07912374001). The strawberry Elongation factor 1 (*EF1*) (Amil-Ruiz *et al*., 2013) gene was used as a control to normalize the amount of cDNA used from each sample. Eurofins PCR Primer Design Tool (https://eurofinsgenomics.eu/en/ecom/tools/pcr-primer-design/) was used to design gene-specific primers for *AP2/EREBP* and *FvHSF* genes (Table S2). Real-time quantitative PCR was carried out using LightCycler 480 SYBR. Green I Master mix (Roche, Cat#04707516001) on a LightCycler® 480 Instrument (F. Hoffmann-La Roche Ltd) with a final volume of 20*μ*l per reaction. Each reaction mixture contained 7ul of Water (PCR grade), 1.0*μ*l cDNA template, 1.0*μ*l of each primer (0.5*μ*M), and 10*μ*l Master Mix (2x). Each reaction was performed in triplicate. Relative gene expression was determined using *EF1* gene as housekeeping gene and analyzed using the qGene protocol of Normalized Expression method (Muller *et al.*, 2002). The primers used for real-time RT-qPCR are listed in Table S2. The data are presented as the mean ± standard error and were tested for statistical significance with an unpaired Student’s t-test. The p < 0.05 was selected as the point of minimal statistical significance in all the analyses.

## Results

### Stress-induced DNA methylation dynamics in *F. vesca*

To evaluate how DNA methylation is altered under diverse plant growth environments, *F. vesca* seedlings were cultivated in a growth chamber *in vitro* for 1 month and then transferred to one of seven different stress environments (Fig. 1a). The treatments represented stress conditions which affect normal plant development (Lämke & Bäurle, 2017): cold, heat, drought, high and low light, and salt as abiotic stresses; salicylic acid (SA) as hormone stress. Control plants were grown on MS medium without stress treatment. After one week of stress exposure, the plants were moved to control conditions for two days to ensure survival. Then, plants were stressed for a second week and finally, moved back to control conditions for two days of recovery (Fig. 1a, see **Materials and Methods** for details). To assess genome-wide DNA methylation levels, DNA was extracted from these plants and submitted to whole genome bisulfite sequencing (WGBS, 20x genome coverage) (Table S1). We carried out a global quantification of DNA methylation in the three sequence contexts (CG, CHG and CHH). First, we combined all bedGraph files of the individual samples into a unionbedg file and filtered the cytosine positions without sequencing coverage and calculated the average DNA methylation levels. Overall, the global DNA methylation levels in all stress conditions were similar (Fig. 1b). *F. vesca* seedlings in control conditions had 40.11% mCG, 14.93% mCHG and 2.38% mCHH. We found substantial decreases of 0.5%, 3.2% and 1.1% for global mCG after cold, heat and salt stress, respectively (Fig. 1b). In addition, we observed a significant global mCHG decrease of 1.8% in salt and an increase of 3.1% in the presence of SA. We did not detect significant changes in global mCHH level. To assess methylation variation in genic and non-genic sequences, we screened the methylome data in all three contexts separately in three regions: 2 kb upstream of genes, along the gene body, and 2 kb downstream of genes within a 100-bp sliding window (Fig. 1c). In contrast to the global analysis, here we observed a higher local DNA methylation variability in the CHH context at the transcription start and end sites (TSS and TES, correspondingly) compared to the other two contexts. Notably for CHH context, cold and heat stress resulted in hypomethylation at the TSS, TES and over the gene body (Fig. 1c). We extracted genes with body methylation (gbM) similarly to the parameters defined previously (Bewick *et al.*, 2019) filtering for at least 20 CGs and a methylation level above the median value. Genes containing gbM showed low variability in all the conditions (Fig. S2).

**Fig. 1.**
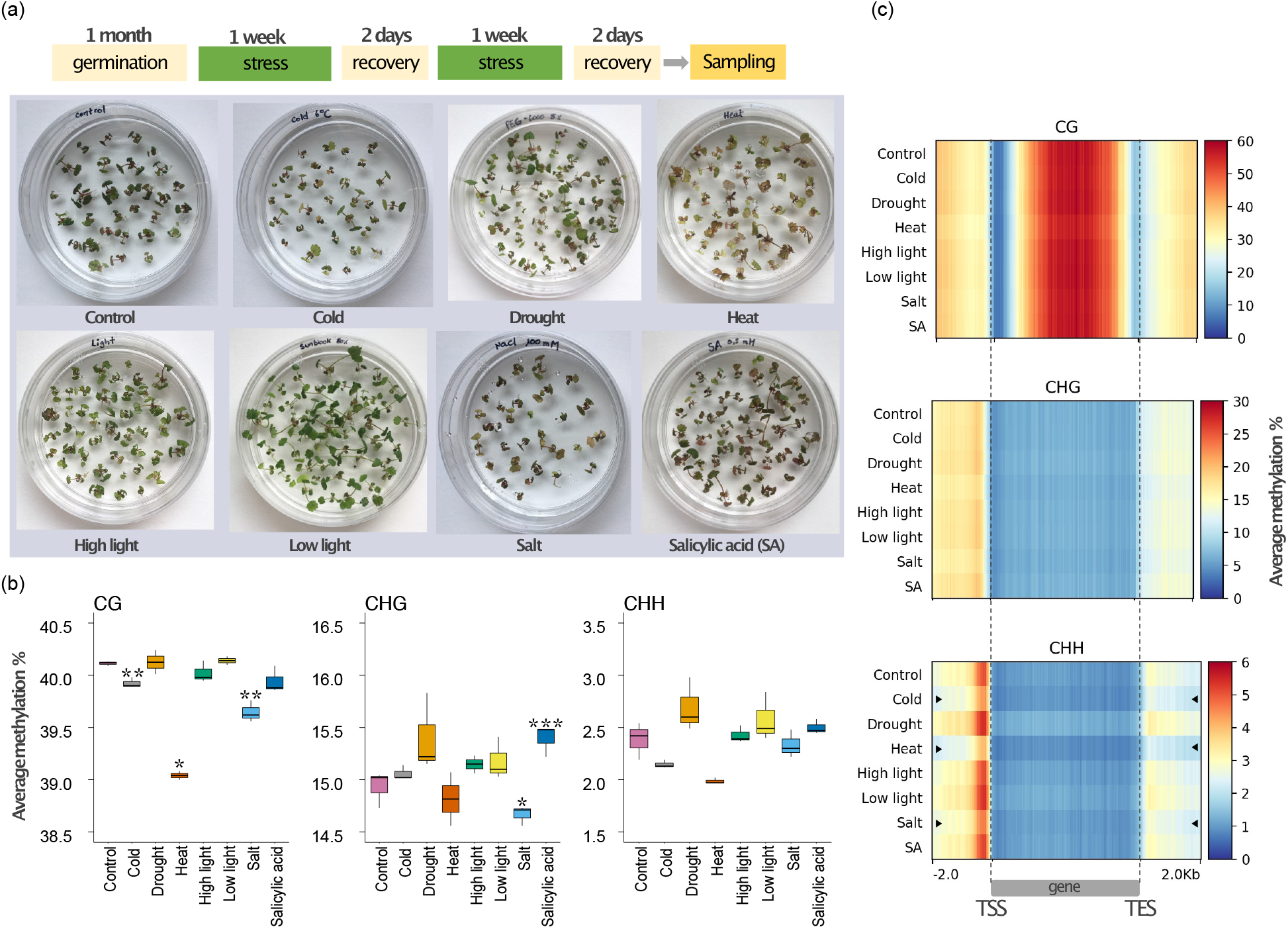
Effect of abiotic and hormone stresses on genome-wide DNA methylation levels in *F. vesca*. (a) Top: Scheme of stress treatment time course. Bottom: Photographs of plates with one-month-old plants grown under different stress conditions (cold, drought, heat, high and low intensity of light, salt, and salicylic acid). (b) Average DNA methylation levels for each cytosine context (CG, CHG, CHH) between normal and stress conditions (only common cytosines positions among all samples were considered that had a minimum coverage of 5 reads). Asterisks indicate levels of significance between treated and control plants: *, p-value < 0.05; **, p-value < 0.01; ***, p-value < 0.001(unpaired two-tailed Student’s t-test). (c) Heat maps showing distribution of DNA methylation (top: CG, middle: CHG, and bottom: CHH context) around genes with and without stress (Control). Mean of the average methylation percentage (within a sliding 100-bp window) was plotted 2 kb upstream of TSS, over the gene body and 2 kb downstream of TES. Black arrows highlight the samples in which a reduction of DNA methylation can be observed in the vicinity of TSS and TES sites.

### Stress particularly affects DNA methylation variability in the non-CG contexts in *F. vesca*

To explore the dynamics of DNA methylation at specific loci in detail, we assessed differentially methylated regions (DMRs) for each sequence context. DMRs were defined using metilene v.0.2.6.1 which uses an algorithm to identify a base-pair window through sequence segmentation with significant methylation differences (Jühling *et al.*, 2016) (See Materials and Methods for parameters). We compared the methylomes of plants submitted to each stress condition with the methylomes from control plants. The majority of DMRs identified were in the CHH context for all conditions, with a maximum of 12,414 DMRs under heat stress. Fewer DMRs were detected in both CG sequence context, ranging from 104 (cold stress) to 3,016 (heat stress), and in the CHG context, ranging from 15 (cold stress) to 236 (heat stress) (Fig. 2a, Table S3). In line with our global DNA methylation analysis, most of the heat, cold and salt stress DMRs were hypomethylated (hypoDMRs) relative to the control condition in all sequence contexts. For drought, high light, low light, and SA stress, most of the DMRs showed hypermethylation (hyperDMRs) in the CHH context (Fig. 2a). Next, to test whether genic and non-genic regions were rich in DMRs, we qualified DMRs on their intersection with promoters (empirically defined as 2 kb to 50 bp upstream TSS), gene bodies, and intergenic regions. The minimum overlap required was 1bp. Many of the CG and CHG DMRs (30-44%, 45-60% respectively) were in genes, while most of CHH DMRs (between 32-44%) were in promoters and intergenic regions (Fig. 2b). In summary, abiotic and hormone stresses led to DNA methylation changes primarily in the CHH context within promoters and intergenic regions. Overall, heat stress caused the most numerous DNA methylation changes.

**Fig. 2.**
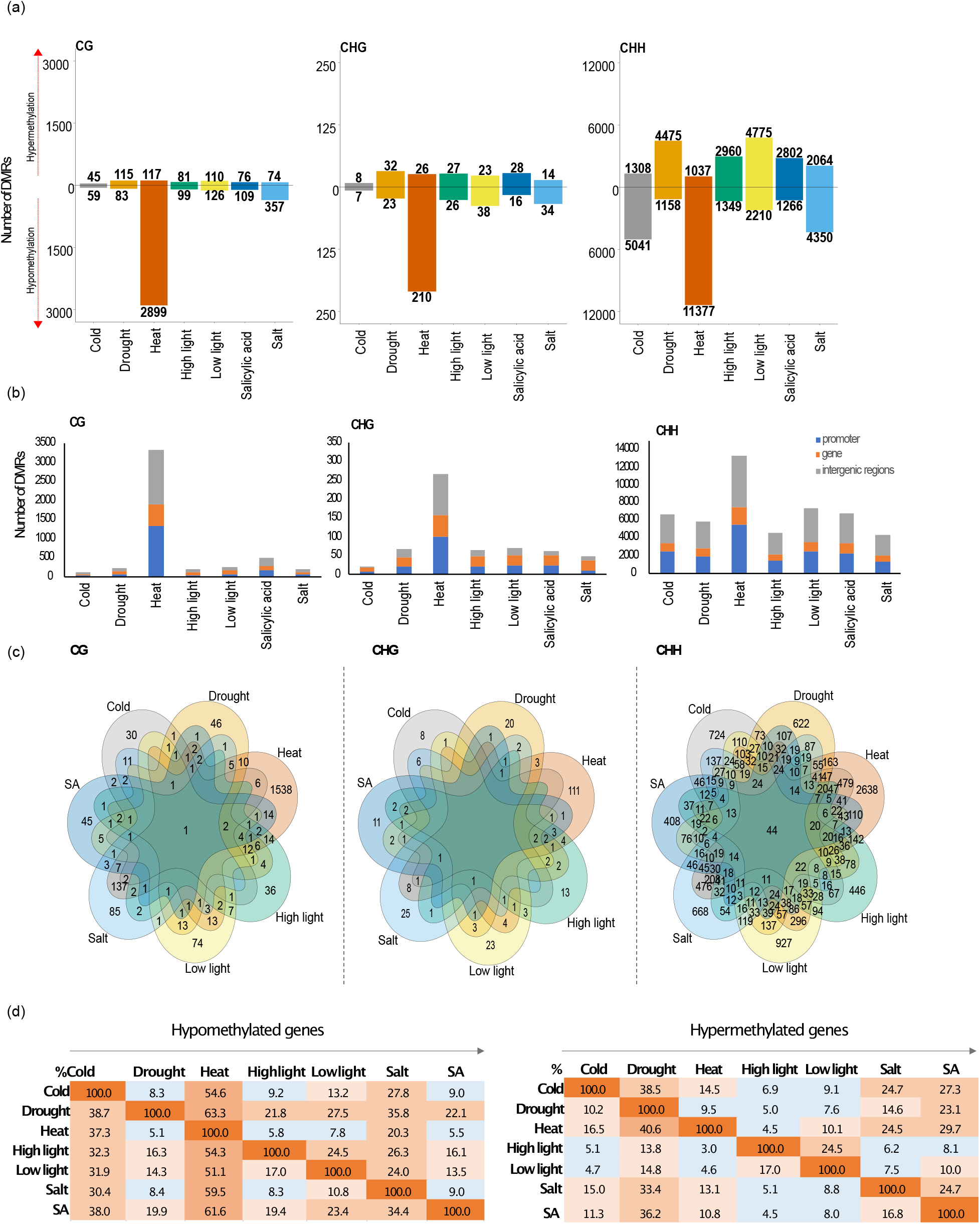
Differentially methylated regions in *F. vesca* seedlings grown at control and stress conditions. (a) Number of stress-induced hyper-(hyperDMRs) and hypomethylated DMRs (hypoDMRs) separated by sequence context. (b) Distribution of DMRs in genomic features: promoter (*2 kb to 50 bp upstream TSS),* gene body, and intergenic regions. Minimum overlap required: 1bp. (c) Venn diagrams of common promoter and genic regions containing hypo- and hyperDMRs per context among all the stress conditions. (d) Percentages of genic loci with hypo- and hyperDMRs which are shared among the stress conditions. Reading from left to the right (arrow). e. g. (Left block) 8.3% genic locations with cold-stressed hypoDMRs overlap with 38.7% genic locations with drought-stress induced hypoDMRs. (Right block) 24% genic loci with hyperDMRs from salt-stressed seedlings overlap with 11% genic loci with hyperDMRs resulting from SA treatment. Ascending intensity of block colors correlate with overlap percentages.

### Identification of stress-induced DNA methylation changes in genic regions

To better understand the potential functional roles of the DMRs and the commonalities between the different stresses, we focused our analysis on DMRs located within promoters and gene bodies (Fig. 2c). We only identified one locus with a CG DMR (within *FvH4_6g40845* an unknown protein) and 44 loci with CHH DMRs that were in common to all stress conditions (Fig. 2c, Fig. S3). Comparing each stress data set of promoter and gene locations with hypo- and hyperDMRs we noted that heat stress shared more loci with hypoDMRs with the other conditions than all other comparisons (Fig. 2d, left). For example, 37.3% of loci with heat-stress hypoDMRs overlapped with 54.6% of all hypoDMRs found under cold-stress. Conversely, drought stress resulted in the highest number of genic loci with hyperDMRs shared with the other conditions. To illustrate, 40.6% of heat-stress hyperDMRs overlapped with 9.5% drought-stress hyperDMRs (Fig. 2d, right). In order to identify potential functional roles for these DMRs, we performed a singular enrichment analysis (SEA) using the AgriGO tool (Tian *et al.*, 2017) (see **Materials and Methods**). *F*. *vesca* has around 34,000 genes but only 54% of which have been assigned a GO number (Li *et al.*, 2019). For this reason, “unknown” annotated genes were omitted for this analysis (Table S4). The analysis was based on the identified genic regions (gene and promoter) with DMRs for each stress condition according to their methylation change (hypo- or hypermethylated). Plants submitted to heat stress showed the largest variation in DNA methylation over genic regions. These were enriched with hyperDMR-associated genes involved in the generation of precursors of metabolites and energy. Genes with heat-stress induced hypoDMRs were enriched for transcription factor (TF) activity, transcriptional regulators, and genes related to cellular components (Table 1). We also found that cold stressed plants had hyperDMR-associated genes enriched for transporter activity (Table 1).

**Table 1.** Gene ontology (GO) enrichment analysis of genic regions (gene and promoter) with DMRs. GO enrichment of genes with hypo- and hyperDMRs caused by thermal stress. The genes are arranged according to their DMRs patterns.

| Stress | Methylation | Description | FDR | Number<br>in input<br>list | Number<br>in<br>BG/Ref | Ontology |
| --- | --- | --- | --- | --- | --- | --- |
| Cold | Hypermethylation | transporter activity | 0.024 | 23 | 838 | F |
| Heat | Hypermethylation | generation of precursor metabolites and energy | 0.021 | 8 | 153 | P |
|  | Hypomethylation | transcription regulator activity | 0.024 | 108 | 420 | F |
|  |  | transcription factor activity | 0.024 | 99 | 380 | F |
|  |  | intracellular | 0.033 | 504 | 2276 | C |
|  |  | cell part | 0.033 | 849 | 3930 | C |
|  |  | cell | 0.033 | 849 | 3930 | C |
|  |  | cytoplasm | 0.033 | 239 | 1029 | C |

GO categories are listed with the false discovery rate-adjusted P-value <0.05. BG: background, Ref: reference, C: Cellular Component; F: Molecular Function; P: Biological process. Only well identified and annotated genes were included in the analysis.

### Heat stress induced hypomethylation at transcription factor coding genes

Transcription factors (TFs) play key roles in plant growth, development, and stress responses. Interestingly, we observed that heat stress resulted in an enrichment of hypoDMRs at promoters and genes related to TFs. Among 99 genes related to transcription factor activity (Table1), 31% were *AP2/EREBP* members and 4% heat shock transcription factors (*HSF*s). *AP2/EREBP* members (119 genes in total) have been characterized in the latest version of the *F. vesca* genome and are known to be involved in stress tolerance (Dong *et al.*, 2021; Xie *et al*., 2019). To evaluate the detailed DNA methylation changes at *AP2/EREBP* genes under different stress conditions, we looked at the distribution of DNA methylation at those loci (Fig. S4a). We observed a noted reduction in DNA methylation at the TSS in the CHH context for heat- and cold-stressed plants. Combining all stress conditions, a total of 74 DMRs were identified within the promoter regions of 44 *AP2/EREBP* genes (Table S5). The marked presence of DMRs within *AP2/EREBP* promoters suggested a relationship between methylation and transcription. We therefore randomly selected four members among each subgroup of the *AP2/EREBP* gene superfamily to detect transcript levels by qRT-PCR analysis (Fig. 3). Even though heat-treated plants seemingly showed higher transcript levels compared to the control plants, only *ERF30* gene was significantly up-regulated (Fig. 3a). In addition, *ERF30* showed the highest methylation difference ratio (CG DMR=-0.42) in its promoter. We also obtained similar results by analyzing the relationship between methylation and expression of heat shock factors (*HSFs).* Initially, 14 *HSFs* have been identified in *F. vesca* genome (Hu *et al.*, 2015); however, in the last genome annotation version (Li *et al.*, 2019), we were able to identify 19 *HSFs* (Table S6). The distribution of methylation over *HSFs* showed high variability in TSS and TES in CHH context compared gene bodies (Fig. S4b). We identified 13 *HSFs* with DMRs within promoter regions mostly attributed to heat stress (Table S6). For the functional expression analysis three genes were randomly selected (Fig. 4). Hypomethylation in promoter regions of *HSFB3a* and *HSFA6a* showed a clear relationship with significantly increased transcript levels after heat stress (Fig. 4a, c). Collectively, our data show that heat stress induces loss of DNA methylation mostly at promoter regions of genes related to specific TF families possibly influencing their transcription.

**Fig. 3.**
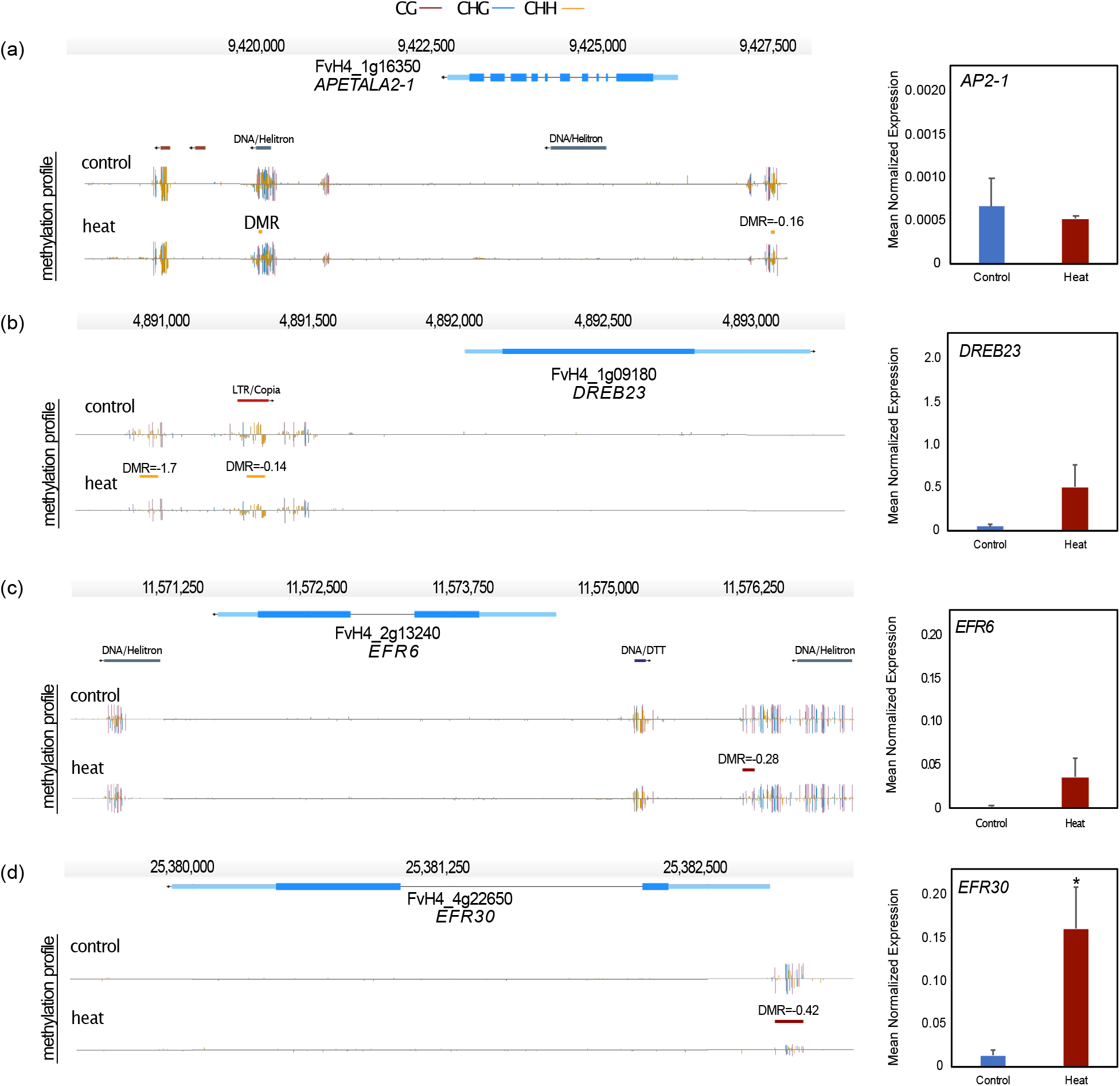
Functional analysis of heat-stress DMRs within *APETALA*2/ethylene-responsive element binding protein (*AP2/EREBP*) superfamily genes. Genome browser views of DMRs located in promoter regions of *AP2/EREBP* genes and bar plots indicate expression ratios using three biological replicates of pooled seedlings. **(a)** *AP2-1 (APETALA2-1),* **(b)** *FvDREB23* (dehydration-responsive element binding 23), **(c)** *ERF6* (ethylene-responsive binding factor 6), (d) *ERF30* (ethylene-responsive binding factor 30). Depicted are genes structures (top panels, UTRs in light blue, exons in blue), TEs (red and dark blue) and DNA methylation levels (histograms). Boxes above the histograms indicate identified DMRs with methylation difference ratios (color codes for DNA methylation: red for CG, blue for CHG and yellow for CHH contexts). Error bars in the plots indicate Standard error. Asterisks indicate levels of significance between treated and non-treated plants: *, p-value < 0.05 (unpaired two-tailed Student’s t-test).

**Fig. 4.**
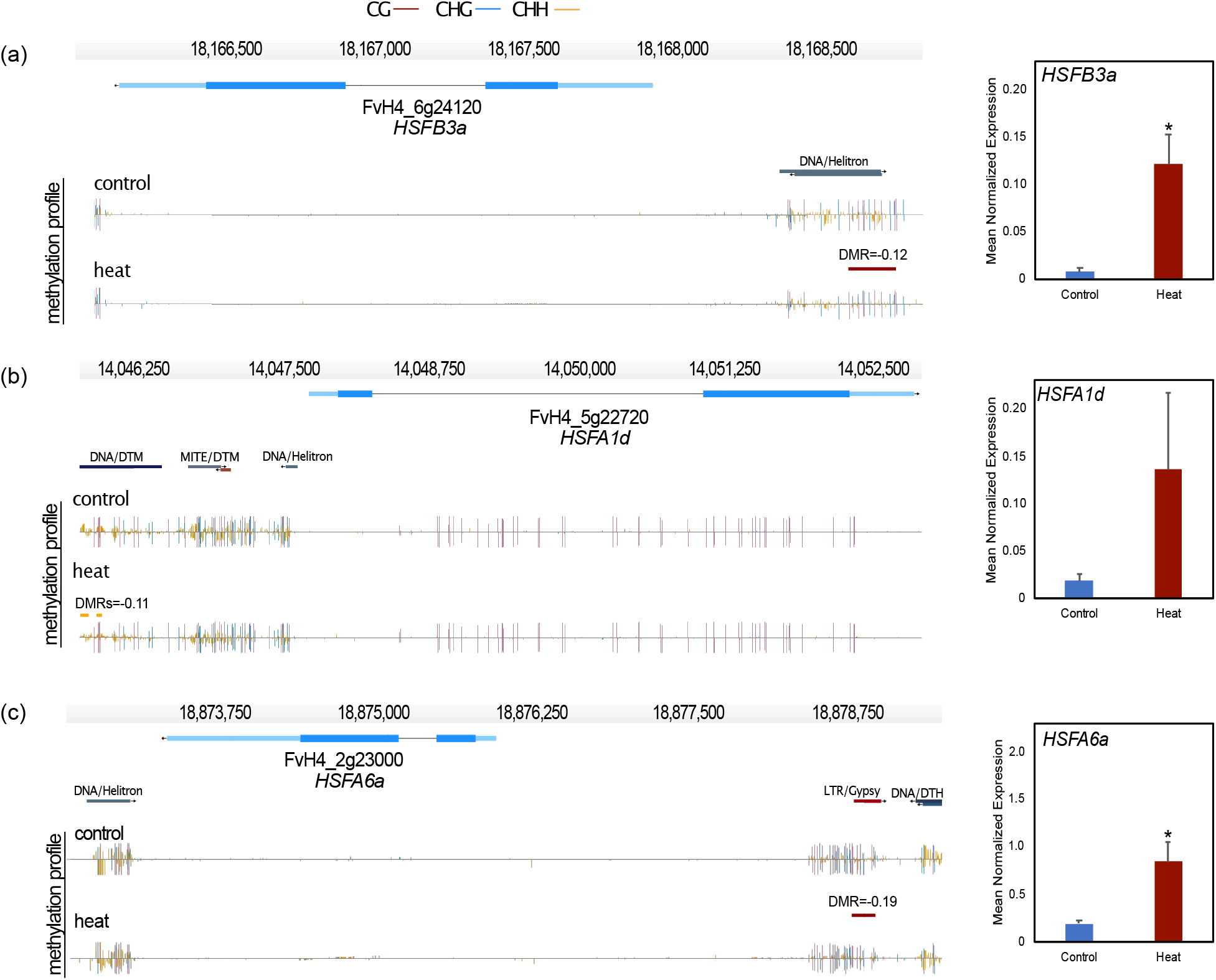
Correlation between heat-stress hypoDMRs and up-regulation of heat shock transcription factors (*HSF*) expression. Genome browser views of DMRs present in located in promoter regions of HSF. (a) *HSFB3a* (heat shock transcription factor B 3a), (b) *HSFA1d* (heat shock transcription factor A 1d), (c) *HSFA6a* (heat shock transcription factor A 6a). Depicted are gene structures (top panels, UTRs in light blue, exons in blue), TEs (red and dark blue) and DNA methylation levels (histograms). Boxes above the histograms indicate identified DMRs with methylation difference ratios (color codes for DNA methylation: red for CG, blue for CHG and yellow for CHH contexts). Error bars in the plots indicate Standard error. Asterisks indicate levels of significance between treated and non-treated plants: *, p-value < 0.05 (unpaired two-tailed Student’s t-test).

### Stress leads to distinct methylation changes at transposable elements (TEs)

Since one of the most important functions of DNA methylation is to repress TE transcription and mobility (Bucher *et al.*, 2012; Deniz *et al.*, 2019; Fedoroff, 2012; Zhang *et al.*, 2018), we investigated the effect of stress on DNA methylation at TEs. To describe the variation in DNA methylation in TE bodies and their flanking regions we plotted DNA methylation profiles in all three contexts. We used 50-bp sliding window: 2kb upstream, over the body and 2 kb downstream (Fig. 5a). In general, all TE families showed high methylation levels in CG context; however, DNA transposons such as the *Mariner* (DTT) and *Helitrons* superfamilies were characterized by low DNA methylation levels in the CHG context. Notably, *Miniature Inverted-Repeat Transposons* (MITEs) showed the highest DNA methylation levels in CHH context. For MITEs, mCHH was distinctly reduced under cold, heat and salt stress (Fig. 5a). To assess significant changes in DNA methylation at specific loci over TEs, we counted the number of DMRs within TE annotations for each stress condition (Fig. 5b). The minimum overlap DMRs/TEs was 1bp. As for genes, we identified the greatest number of DMRs within TEs in heat-stressed plants in all cytosine contexts (Fig. 5b). Cold, heat, and salt stress displayed more hypoDMRs in TEs compared to drought, low light, high light, and SA, which produced more hyperDMRs in TEs (Fig. 5c). Although there was no specific TE family significantly enriched in DMRs, heat-stress resulted in at least 12% of all MITES acquiring hypoDMRs (Fig. 5c, Table S7) and most of them were close to genic regions (< 2kb upstream genes) (see Fig. 6 for examples). Overall, these results suggested that DNA methylation dynamics among all TE members changed proportionally in all superfamilies; nonetheless, the change regarding gain or loss of DNA methylation was clearly defined by the applied stresses.

**Fig. 5.**
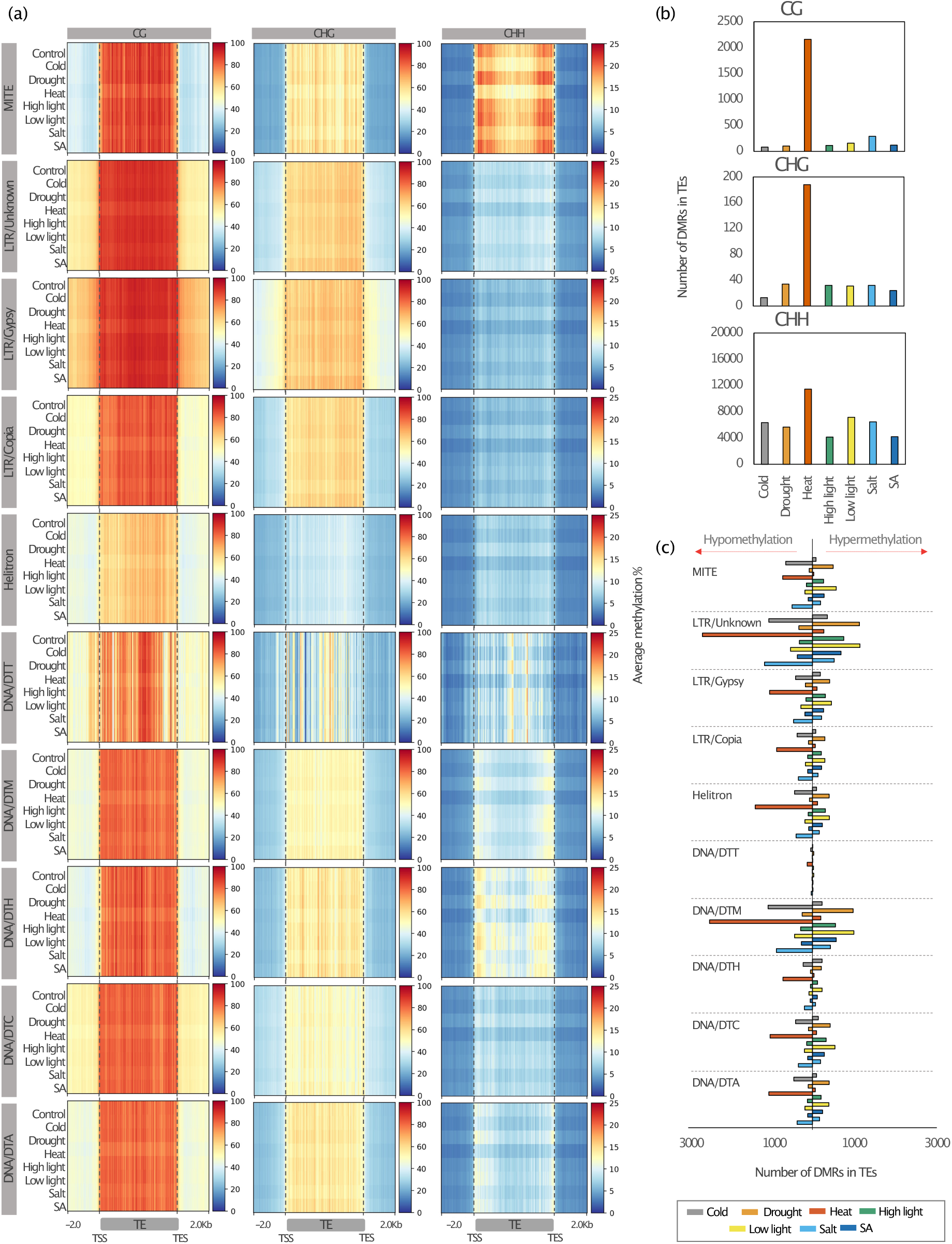
Association of stress-induced differentially methylated regions with transposable elements in *F. vesca.* (a) Heatmaps showing DNA methylation profiles for the all the TE families separated by sequence context mCG (left), mCHG (center) and mCHH (right). The mean of the average DNA methylation percentage (within a 50 bp sliding window) was plotted for the TE bodies and 2 kb around the TSS and TES regions. (b) Number of stress induced DMRs in TEs per sequence context. (c) Number of hypoDMRs (left) and hyperDMRs (right) within different TE families. Class I elements (retrotransposons): *LTR-Copia*, *LTR-Gypsy*. Class II elements (DNA transposons): TIR: *Tc1–Mariner* (DTT), *hAT* (DTA), *Mutator* (DTM), *PIF– Harbinger* (DTH), *CACTA* (DTC); *Helitron; Miniature Inverted-Repeat Transposons* (MITEs). Upper char box indicates the colors which represent each treatment.

**Fig. 6.**
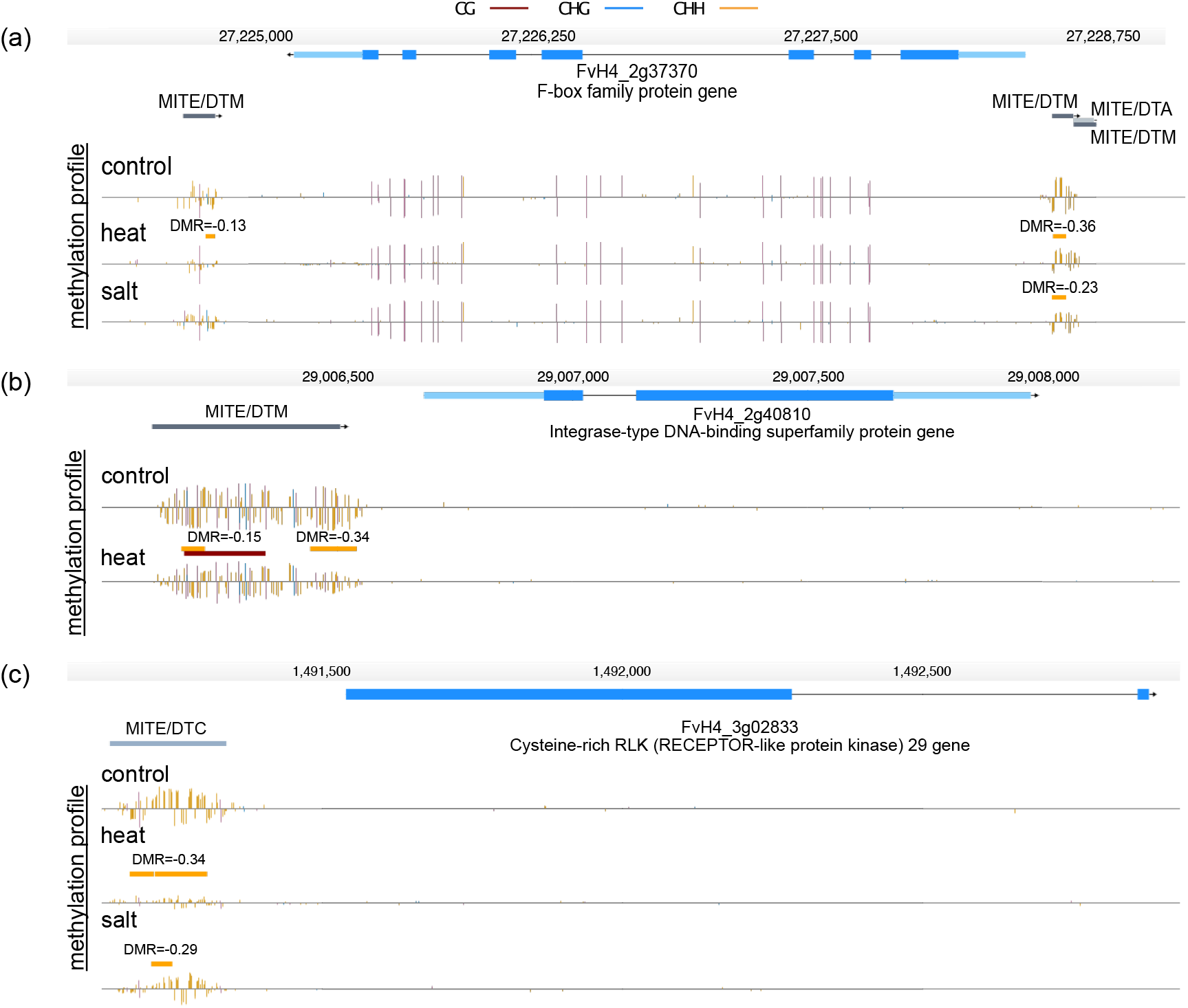
*Miniature Inverted-Repeat Transposons* (MITEs) present hypomethylated regions (DMRs) under abiotic stress conditions. Genome browser views of DMRs present in MITEs located near genes. (a) F-box family protein gene (FvH4_2g37370). (b) Integrase-type DNA binding superfamily protein gene (FvH4_2g40810). (c) Cysteine-rich RLK (RECEPTOR-like protein kinase) 29 (FvH4_3g02833). Depicted are genes structures (top panels, UTRs in light blue, exons in blue), TEs (red and dark blue) and DNA methylation levels (histograms). Boxes above the histograms indicate identified DMRs with methylation difference ratio (color codes for DNA methylation: red for CG, blue for CHG and yellow for CHH contexts).

### DMRs are enriched in distinct regions of the *F. vesca* in the genome

Even though genome-wide DNA methylation variation levels were low (Fig. 1b), we observed regions in the genome that were enriched in DMRs in all stress conditions (Fig. 7a). Indeed, we observed DMR hotspots in the *F. vesca* genome. Notably, the distribution showed a similar pattern on all chromosomes independently of the stress conditions. The DMR density was not clearly related to gene and TE density (Fig. 7b); nevertheless, when we analyzed individual TE families, *Helitron* density hotspots showed a clear correlation with DMR density (Fig. 7). Currently, little is known about the exact localization of centromeric and pericentromeric regions in strawberry (Li *et al.*, 2018). However, it has been reported previously that centromeric regions can be enriched in *Helitrons* in certain plant genomes (Xiong *et al.*, 2016). It is important to note here, that the *Helitrons* themselves were not enriched in DMRs.

**Fig. 7.**
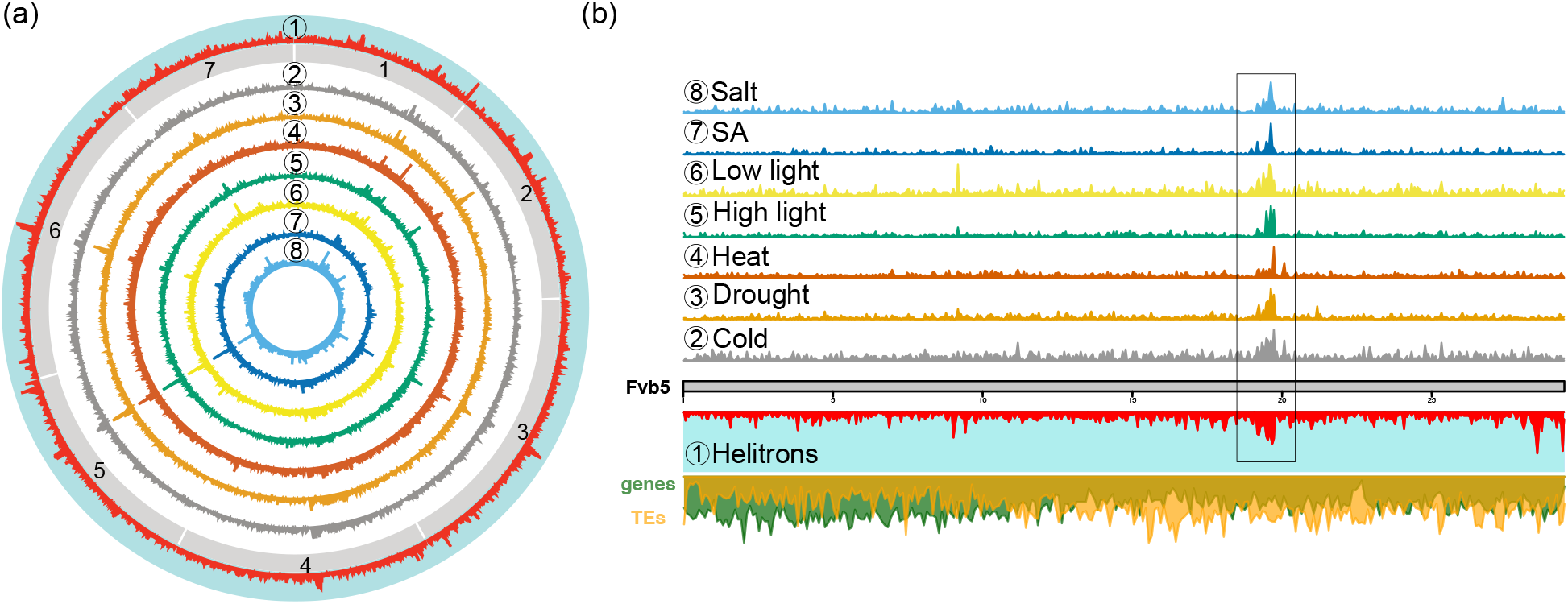
Genome-Wide distribution of DMR densities. (a) Circa plot showing DMR densities on all 7 strawberry chromosomes (gray boxes) for each stress condition (2. Cold, 3. Drought, 4. Heat, 5. High light, 6. Low light, 7. SA, and 8. Salt). *Helitron* density (1) in the genome is depicted on the outer most circle in red with turquoise background. (b) DMR density depicted on chromosome 5 (Fvb5) displaying a common enrichment among all the stress conditions (left of the 20 Mb tick mark). Below, the chromosome is indicated in grey, green shows gene density, yellow the TE density and red Helitrons along the chromosome.

## Discussion

### Genome-wide DNA methylation patterns are altered under stressful environmental conditions

Environmentally induced epigenetic changes include mechanisms that respond to abiotic or biotic stress conditions in plants (Verhoeven *et al.*, 2016). DNA methylation variation might trigger modifications in plant development and physiology contributing to phenotypic variability, thereby presumably contributing to plant acclimation (Dubin *et al.*, 2015; Lämke & Bäurle, 2017; Xu *et al.*, 2020; Zhang *et al.*, 2021). Here, we studied DNA methylation dynamics in *F. vesca* submitted to two successive stress applications. Looking at the total DNA methylation levels in the three different sequence contexts, we observed only slight variations among the different stress conditions. However, by performing a DMR analysis we detected extensive differences at specific loci especially in the CHH context. This is in agreement with studies which reported that mCHH was most dynamic in response to different climates (Dowen *et al.*, 2012; Dubin *et al.*, 2015; Kenchanmane *et al.*, 2019). More specifically, in the case of *F. vesca,* altitude variations of natural strawberry populations was found to be correlated with a high variability in DNA methylation in the CHH context (De Kort *et al.*, 2020; Kort *et al.*, 2021). In our study, cold, heat, and salt stress resulted in substantial local loss of DNA methylation in all sequence contexts, particularly in regions close to TSS and TES sites in the *F. vesca* genome (Fig. 1c). Such DNA methylation changes have been associated with different degrees of cold tolerance. Indeed, different crop species such as maize, rice, cotton, and chickpea presented an increased number of hypomethylated regions near abiotic stress response genes and transcription factors upon cold stress (Fan *et al.*, 2013; Pan *et al.*, 2011; Rakei *et al.*, 2016; Shan *et al.*, 2013). Higher temperature resulted in hypomethylation in different plant tissues, such as rice seeds, soybean roots, and tobacco leaves, affecting plant growth by altering gene expression patters in genes that control biosynthesis and catabolism of phytohormones (Centomani *et al*., 2015; Hossain *et al.*, 2017; Qian *et al.*, 2019; Suriyasak *et al.*, 2021). In addition, salt stress induced epigenetic variation in Arabidopsis have been shown to be partially transmitted to offspring, primarily via the female germ line (Wibowo *et al*., 2016). On the other hand, we noticed little gain of global DNA methylation caused by different intensities of light or drought and SA stresses; yet causing a considerable number of hyperDMRs in the CHH context. Different results were found in tomato plants exposed to variable intensities of light which resulted in DNA hypomethylation and transcriptional changes causing male-sterility (Omidvar & Fellner, 2015). However, drought stress induced mCG and mCHG hypermethylation and a slight decrease in mCHH methylation in mulberry (Li *et al.*, 2020). There is also evidence that drought, nitrogen-deficiency and heavy metal stresses can result in heritable changes in DNA methylation levels across generations in rice (Kou *et al*., 2011; Ou *et al.*, 2012; Zheng *et al.*, 2013). Another study showed that DNA methylation changes induced by hormone stresses via Jasmonic and Salicylic acid can be faithfully transmitted to offspring by asexual reproduction in dandelions (Verhoeven *et al.*, 2010). Together, these findings open the question of whether stress-induced DNA methylation changes found in *F. vesca* can be maintain over generations through asexual (stolons) and sexual (seeds) reproduction. An aspect we are currently intensively investigating.

### Different TE superfamilies show contrasted responses to stresses

DNA methylation plays a key role in limiting transcriptional activation and mobilization of TEs in order to ensure genome integrity (Slotkin & Martienssen, 2007; Bucher *et al.*, 2012; Deniz *et al.*, 2019). Indeed, TEs can be an important source of genetic and epigenetic variation that can influence stress-responses (Naito *et al*., 2009). Here, we wanted to better understand how different TE superfamilies respond to the stresses we applied. Using this approach, we observed distinct DNA methylation profiles at TEs depending on their superfamily and the applied stress conditions. Overall, we found that *F. vesca* TEs are highly methylated in both, CG and CHG sequence contexts. Similar results were observed in maize where LTR and TIR elements are highly methylated under normal conditions (Noshay *et al.*, 2019). On the other hand, we noticed that overall DNA methylation levels in the CHH context were lower and more variable among different TE superfamilies and stress conditions, implying contrasted responses of TEs to stresses and that these TEs are silenced by different transcriptional gene silencing pathways. These DNA methylation changes could have direct physiological impacts as non-CG methylation seems to create a boundary between genes and TEs (Kenchanmane *et al*., 2019). For example, it has been found that TEs located close to stress-induced genes in Arabidopsis and rice are silenced by hypermethylation after phosphate starvation in order to prevent collateral activation of TE transcription during stress (Secco *et al.*, 2015). In our study we found that heat stress affected mCHH in the TE body of all TE superfamilies. MITES had the highest percentage of members targeted by DMRs (Table S7). In addition, we noticed MITES to often be present in promoter regions of genes with DMRs. A genome-wide transcriptional analysis would be needed to generally conclude on whether the variation of DNA methylation in promoters might be stimulated by the presence of MITES thereby contributing to gene expression control *in cis*. In line with these observations, previous studies have highlighted the importance of MITES in genome evolution and how MITE insertions in promoter regions can regulate the expression of genes in a wide variety plant species such as maize, rice, and mulberry (Lu *et al*., 2012; Wei *et al.*, 2014; Mao *et al*., 2015; Xin *et al*., 2019). Taken together, these observations suggest that specific TE superfamily members with dynamic DNA methylation levels may contribute to stress response strategies in plants.

### DMR location preferences for centromeric regions

It has long been established that DNA methylation is enriched in peri-/centromeric regions which follow the distribution of TEs over the chromosomes of genomes of the Brassicaceae family, as recently also confirmed for *Thlaspi arvense* (Seymour *et al.*, 2014; X. Zhang *et al.*, 2006; Naish *et al.*, 2021; Nunn *et al.*, 2022;). In our study, investigating the global distribution of DMRs over the *F. vesca* chromosomes, we found that regions with high DMR density correlated with regions enriched in *Helitrons*. Interestingly, we made this observation for all stress conditions (Fig. 7a, b). Currently, *F. vesca* centromeres are not well defined; however, a genome-wide scan of the *F. vesca* genome for tandem repeats suggested the presence of *Helitrons* near the centromeres (Xiong *et al*., 2016). This suggests that *F. vesca* centromeres or pericentromeric regions respond to stresses with DNA methylation changes. To further confirm the exact localization of *F. vesca* centromeres, immunoprecipitations using a CENH3 antibody followed by sequencing will be required (Comai *et al.*, 2017). Why centromere-associated regions are more prone to DNA methylation changes and the physiological relevance of this observation still needs to be determined.

### Heat-stress results in loss of methylation in regions flanking TFs

In our study, the global analysis of the DMRs resulting from each stress condition highlighted numerous shared genomic patterns among the stresses but also interesting stress-specific genomic features. In the case of heat stress, we observed hypoDMRs predominantly in gene, promoter, and TE regions. Extreme temperatures are one of the main stresses affecting plants that particularly alter their development and potentially cause yield loss (Janni *et al.*, 2020). Heat response triggers a chain of highly conserved mechanisms ( Ohama *et al.*, 2017; de Vries *et al.*, 2020; Medina *et al.*, 2021). In the case of *F. vesca,* we found that DNA methylation differences were enriched in genes related to transcription factor regulation and activity as well as generators of metabolites and energy. This result is consistent with the idea that transcription factors are required to reprogram stress-related genes (Ohama *et al.*, 2016). There is evidence which showed heat stress response requires not only transcription factor activity but also epigenetic regulators and small RNAs to rapidly activate genes (Ohama *et al*., 2017). However, little is known about how these responses are influenced directly by changes in DNA methylation or *vice versa*. Transcription factor families such as *AP2/EREBP* and *HSFs* play important roles in response to abiotic stresses in plants including strawberry, apple and maize (Brown *et al.*, 2016; H. C. Liu *et al.*, 2011; Qian *et al.*, 2019; Xie *et a*l., 2019; C. L. Zhang *et al.*, 2020). More over, some *AP2/EREBP* genes are known to be highly induced under heat stress conditions and up-regulated by HSFs through an interconnected stress regulatory network (Q. Liu *et al.*, 1998; H. C. Liu *et al.*, 2011). Here, we provide epigenetic evidence which suggests that members of the *AP2/EREBP* might be regulated by changes in DNA methylation in *F. vesca*. Among them, the promoter region of the ethylene response factor *EFR30* gene showed loss of methylation and significant up-regulation after heat stress. Recent studies showed that ERFs enhance basal thermotolerance by regulating heat-responsive genes and interacting with HSF in Arabidopsis and tomato (Klay et al., 2018; Huang *et al.*, 2021). Similarly, hypomethylation in the TSS of genes involved in the control of cell growth in tobacco and stress-tolerance genes in maize after heat stress exposure is consistent with the increase in their transcription levels (Centomani *et al*., 2015; Qian *et al.*, 2019). Correspondingly, we identified 58% of *HSFs* genes with hypoDMRs in their promoter regions after heat stress. *HSFs* are crucial for thermotolerance capacity and regulate the expression of several heat-stress response genes as Heat shock proteins (HSPs) (Liu *et al.*, 2011). Here, we showed up-regulation of class A and B *HFS*s in *F. vesca* after heat stress. Comparable results were obtained in a transcriptome analysis of the octoploid strawberry where *HSF* expression was induced by a heat shock treatment (Liao *et al.*, 2016). Taken together, these findings provide insights into stress induced DNA methylation as plant response, and it will be of great interest to investigate the role of DNA hypomethylation in promoter regions of *AP2/EREBP* and *HSF* genes in regulating or priming transcription during heat stress in *F. vesca*.

## Conclusions

In summary, our data revealed how DNA methylation profiles at genes and transposable elements can vary in response to stresses in wild strawberry. In addition, we observed correlations between changes in DNA methylation and gene regulation as interconnected mechanisms during stress exposure. We provide insights into how specific chromosomal regions can vary at DNA methylation levels under stress conditions. These observations suggest that the epigenetic flexibility of centromeres may play an important role during plant stress response. Furthermore, using *F. vesca* as a model plant will help to better understand the stress response of more complex genomes in the Rosacea family. Overall, this study with high-resolution methylome mapping of the *F. vesca* genome will contribute to a better comprehension of epigenetic responses under variable growth conditions. It remains to be tested if such epigenetic changes can be inherited during sexual or clonal propagation (which is common in *F. vesca*) and if such changes could contribute to adaptation to changing environments.

## Acknowledgements

This study was supported by INRAE, Angers-Nantes, France and Agroscope, Nyon-Switzerland. We would like to thank all the members of the EpiDiverse consortium (www.epidiverse.eu) and the Crop Genome Dynamics research group for invaluable support, Katharina Jandrasits for preparing WGBS libraries and Dr. Marta Robertson for the careful reading of the manuscript.

## Author Contribution

ME.L and E.B conceived the study. ME.L performed the experiments, analyzed the methylome data and wrote the manuscript. D.R. assembled the *F. vesca* genome and wrote the manuscript. C.B. performed experiments and wrote the manuscript. B.D wrote the manuscript. E.B. designed experiments, analyzed data, set up the genome browser and wrote the manuscript.

