## Supporting Information for "DNA methylation dynamics during stress-response in woodland strawberry (*Fragaria vesca*)"

Additional supporting information may be found in the online version of this article

**Table S1**. Bisulfite sequencing data quality analysis
**Table S2.** Primers used for real-time RT-qPCR
**Table S3**. Summary of Differentially methylated regions (DMRs) in strawberry seedling grown at normal and stress conditions
**Table S4**. Percentage of genes used for GO enrichment analysis
**Table S5.** APETALA2/ethylene-responsive element binding protein (AP2/EREBP) superfamily which present DMRs in all different contexts
**Table S6**. Heat shock transcription factors *(HSFs)* which contains DMRs under stress
**Table S7**. Association of stress-induced differentially methylated regions with transposable elements in F. vesca. (a) Total number of TE in the *Fragaria vesca* genome. (b) Number of TEs with differentially methylated regions.

**Fig. S1 Heatmaps of significant DMRs (q < 0.05) in CG, CHG and CHH**. Methylome comparisons from control plants vs a stress condition: (a) cold, (b) drought, (c) heat, (d) high light, (e) low light, (f) SA, (g) salt.
**Fig. S2 Global methylation plot profiles of genes with body methylation (gbM).** Plots show distribution of DNA methylation in CG, CHG and CHH context around genes classified as genes with body methylation (gbM) with and without stress (Control). Mean of the average methylation percentage (within a sliding 100-bp window) was plotted 2 kb upstream of TSS, over the gene body and 2 kb downstream of TES.
**Fig. S3 Common stress induced DMRs per context in promoter and genic regions.** (a) Overlapping CG-DMRs with different methylation profiles in all stress conditions. (b) CHG-DMRs diverge in different stress conditions and are enriched in gene bodies. Genome browser view methylation profiles derived from whole genome bisulfite sequencing. Boxes above the histograms indicate identified DMRs (color codes for DNA methylation: red for CG, blue for CHG and yellow for CHH contexts).

**Fig. S4 Global CHH methylation distribution over transcription factors under abiotic and hormone stress conditions**. (a) APETALA2/ethylene-responsive element binding protein (*AP2/EREBP*) superfamily. (b) Heat shock transcription factors *(HSF)*. Heatmaps showing distribution of DNA methylation in CHH context around genes with and without stress (Control). Mean of the average methylation percentage (within a sliding 100-bp window) was plotted 2 kb upstream of TSS, over the gene body and 2 kb downstream of TES.

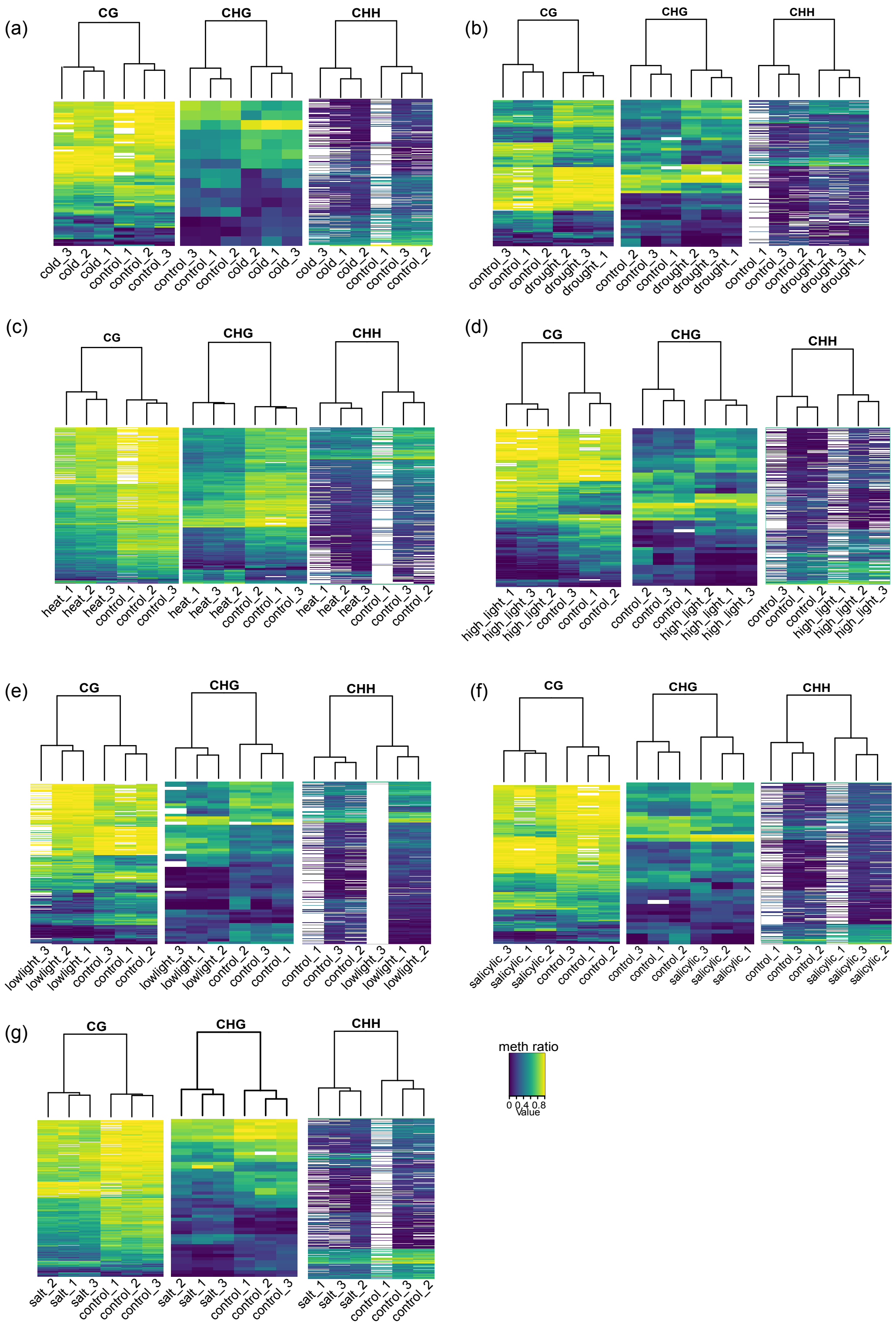


**Fig. S1 Heatmaps of significant DMRs (q < 0.05) in CG, CHG and CHH**. Methylome comparisons from control plants vs a stress condition: (a) cold, (b) drought, (c) heat, (d) high light, (e) low light, (f) SA, (g) salt.

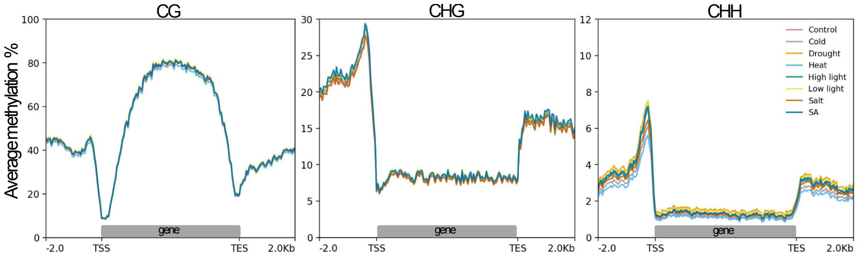


**Fig. S2 Global methylation plot profiles of genes with body methylation (gbM).** Plots show distribution of DNA methylation in CG, CHG and CHH context around genes classified as genes with body methylation (gbM) with and without stress (Control). Mean of the average methylation percentage (within a sliding 100-bp window) was plotted 2 kb upstream of TSS, over the gene body and 2 kb downstream of TES.

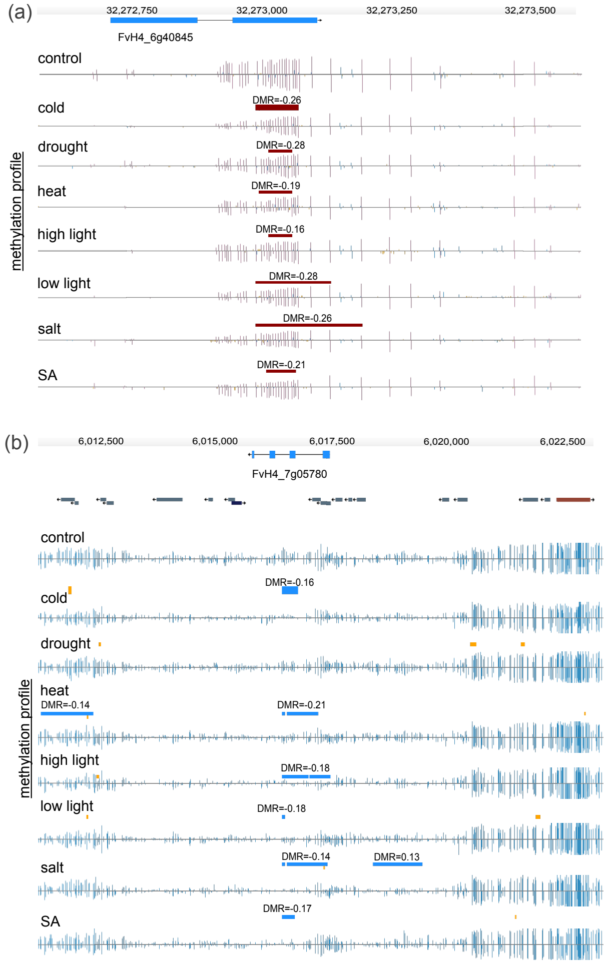


**Fig. S3 Common stress induced DMRs per context in promoter and genic regions.** (a) Overlapping CG-DMRs with different methylation profiles in all stress conditions. (b) CHG-DMRs diverge in different stress conditions and are enriched in gene bodies. Genome browser view methylation profiles derived from whole genome bisulfite sequencing. Boxes above the histograms indicate identified DMRs (color codes for DNA methylation: red for CG, blue for CHG and yellow for CHH contexts).

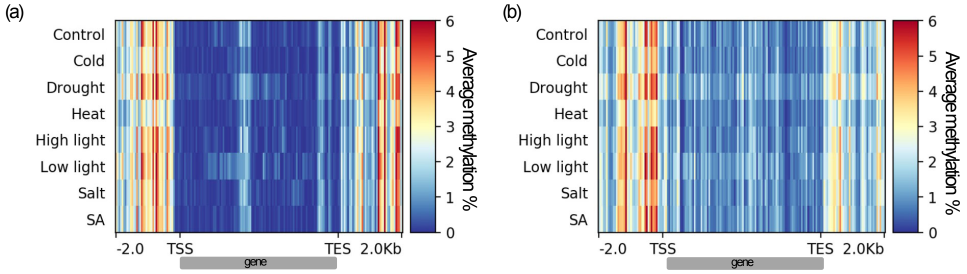


**Fig. S4 Global CHH methylation distribution over transcription factors under abiotic and hormone stress conditions**. (a) APETALA2/ethylene-responsive element binding protein (*AP2/EREBP*) superfamily. (b) Heat shock transcription factors *(HSF)*. Heatmaps showing distribution of DNA methylation in CHH context around genes with and without stress (Control). Mean of the average methylation percentage (within a sliding 100-bp window) was plotted 2 kb upstream of TSS, over the gene body and 2 kb downstream of TES.

**Table S1.** Bisulfite sequencing data quality analysis

| Sample | Treatment | Tissue | Tech.rep. | strategy | source | selection | layout | platform | Model | Total raw sequences | reads mapped | % | reads mapped and paired | properly paired reads (%) | Non-conversion Rate |
|---|---|---|---|---|---|---|---|---|---|---|---|---|---|---|---|
| FV_FR_01_01_P0_IC0_S1 | control | Seedling | 1 | Bisulfite-Seq | GENOMIC | RANDOM | paired | ILLUMINA | HiSeq X Ten | 54350112 | 43194568 | 79.47 | 41103610 | 89.6 | 0.0849765 |
| FV_FR_01_01_P0_IC0_S2 | control | Seedling | 1 | Bisulfite-Seq | GENOMIC | RANDOM | paired | ILLUMINA | HiSeq X Ten | 75685306 | 61502655 | 81.26 | 59033115 | 88.7 | 0.0957948 |
| FV_FR_01_01_P0_IC0_S3 | control | Seedling | 1 | Bisulfite-Seq | GENOMIC | RANDOM | paired | ILLUMINA | HiSeq X Ten | 88196726 | 71859155 | 81.47 | 68666898 | 88.5 | 0.0824866 |
| FV_FR_01_01_P0_IT0_S1 | heat | Seedling | 1 | Bisulfite-Seq | GENOMIC | RANDOM | paired | ILLUMINA | HiSeq X Ten | 82033770 | 63336642 | 77.20 | 60355952 | 87.1 | 0.158832 |
| FV_FR_01_01_P0_IT0_S2 | heat | Seedling | 1 | Bisulfite-Seq | GENOMIC | RANDOM | paired | ILLUMINA | HiSeq X Ten | 95891176 | 78466389 | 81.82 | 76205355 | 90.1 | 0.0658496 |
| FV_FR_01_01_P0_IT0_S3 | heat | Seedling | 1 | Bisulfite-Seq | GENOMIC | RANDOM | paired | ILLUMINA | HiSeq X Ten | 131601020 | 107575452 | 81.74 | 104747479 | 89.6 | 0.0489135 |
| FV_FR_01_01_P0_IF0_S1 | cold | Seedling | 1 | Bisulfite-Seq | GENOMIC | RANDOM | paired | ILLUMINA | HiSeq X Ten | 75356018 | 62062067 | 82.35 | 60460228 | 89.9 | 0.0557463 |
| FV_FR_01_01_P0_IF0_S2 | cold | Seedling | 1 | Bisulfite-Seq | GENOMIC | RANDOM | paired | ILLUMINA | HiSeq X Ten | 113833012 | 92894326 | 81.60 | 90539656 | 89.5 | 0.0523164 |
| FV_FR_01_01_P0_IF0_S3 | cold | Seedling | 1 | Bisulfite-Seq | GENOMIC | RANDOM | paired | ILLUMINA | HiSeq X Ten | 62087752 | 51227464 | 82.50 | 49991019 | 89.3 | 0.0709958 |
| FV_FR_01_01_P0_IN0_S1 | salt | Seedling | 1 | Bisulfite-Seq | GENOMIC | RANDOM | paired | ILLUMINA | HiSeq X Ten | 65068602 | 54636225 | 83.96 | 53200844 | 91.2 | 0.113733 |
| FV_FR_01_01_P0_IN0_S2 | salt | Seedling | 1 | Bisulfite-Seq | GENOMIC | RANDOM | paired | ILLUMINA | HiSeq X Ten | 100511800 | 84669826 | 84.23 | 82590659 | 90 | 0.0612594 |
| FV_FR_01_01_P0_IN0_S3 | salt | Seedling | 1 | Bisulfite-Seq | GENOMIC | RANDOM | paired | ILLUMINA | HiSeq X Ten | 80760776 | 67889432 | 84.06 | 66159488 | 89.7 | 0.0599842 |
| FV_FR_01_01_P0_IS0_S1 | SA | Seedling | 1 | Bisulfite-Seq | GENOMIC | RANDOM | paired | ILLUMINA | HiSeq X Ten | 57969684 | 47564328 | 82.05 | 46310855 | 89.6 | 0.0529798 |
| FV_FR_01_01_P0_IS0_S2 | SA | Seedling | 1 | Bisulfite-Seq | GENOMIC | RANDOM | paired | ILLUMINA | HiSeq X Ten | 91659264 | 71433867 | 77.93 | 69656974 | 90.4 | 0.062181 |
| FV_FR_01_01_P0_IS0_S3 | SA | Seedling | 1 | Bisulfite-Seq | GENOMIC | RANDOM | paired | ILLUMINA | HiSeq X Ten | 117736938 | 96487769 | 81.95 | 93542164 | 89 | 0.0828464 |
| FV_FR_01_01_P0_IB0_S1 | low light | Seedling | 1 | Bisulfite-Seq | GENOMIC | RANDOM | paired | ILLUMINA | HiSeq X Ten | 85964910 | 71004318 | 82.59 | 68857406 | 90 | 0.0754448 |
| FV_FR_01_01_P0_IB0_S2 | low light | Seedling | 1 | Bisulfite-Seq | GENOMIC | RANDOM | paired | ILLUMINA | HiSeq X Ten | 81716372 | 66908386 | 81.87 | 64790729 | 89.8 | 0.0829156 |
| FV_FR_01_01_P0_IB0_S3 | low light | Seedling | 1 | Bisulfite-Seq | GENOMIC | RANDOM | paired | ILLUMINA | HiSeq X Ten | 29863420 | 24419321 | 81.77 | 23741172 | 89.9 | 0.232153 |
| FV_FR_01_01_P0_ID0_S1 | drought | Seedling | 1 | Bisulfite-Seq | GENOMIC | RANDOM | paired | ILLUMINA | HiSeq X Ten | 109809600 | 91100946 | 82.96 | 88787544 | 89.9 | 0.109024 |
| FV_FR_01_01_P0_ID0_S2 | drought | Seedling | 1 | Bisulfite-Seq | GENOMIC | RANDOM | paired | ILLUMINA | HiSeq X Ten | 94456118 | 77908248 | 82.48 | 75691073 | 85.3 | 0.301095 |
| FV_FR_01_01_P0_ID0_S3 | drought | Seedling | 1 | Bisulfite-Seq | GENOMIC | RANDOM | paired | ILLUMINA | HiSeq X Ten | 97889484 | 80948114 | 82.69 | 78542173 | 89.1 | 0.0902901 |
| FV_FR_01_01_P0_IL0_S1 | high light | Seedling | 1 | Bisulfite-Seq | GENOMIC | RANDOM | paired | ILLUMINA | HiSeq X Ten | 54431008 | 44156034 | 81.12 | 42787879 | 88.7 | 0.0792454 |
| FV_FR_01_01_P0_IL0_S2 | high light | Seedling | 1 | Bisulfite-Seq | GENOMIC | RANDOM | paired | ILLUMINA | HiSeq X Ten | 79622196 | 64766856 | 81.34 | 62804109 | 89.6 | 0.0868833 |
| FV_FR_01_01_P0_IL0_S3 | high light | Seedling | 1 | Bisulfite-Seq | GENOMIC | RANDOM | paired | ILLUMINA | HiSeq X Ten | 60025776 | 48357441 | 80.56 | 46943047 | 90.2 | 0.0792454 |

**Table S2.** Primers used for real-time RT-qPCR

| Name | Primer | Gene ID |
|---|---|---|
| FvEF1a_forward | CCATGGTTGTTGAAACTTTCTC | FvH4_7g20050 |
| FvEF1a_reverse | GGC GCA TGT CCC TCA CAG | |
| EFR30_forward | TTTCAATACCACCCTCTCCC | FVH4_4g22650 |
| EFR30_reverse | GTAACTCTACTCGGCAAAGATG | |
| EFR6_forward | GAGATAGAAACAAAGCGGCAC | FVH4_2g13240 |
| EFR6_reverse | CAC CAC CGA GCT GAA AAA ATA ATA | |
| AP2_forward | AAA AGA GAG GAG AGA GAG AGG | FvH4_1g16350 |
| AP2_reverse | ACT CTA CGA GAG AAA ACA AAG AAC | |
| DREB23_forward | CTCCACACCCAAACACATC | FVH4_1g09180 |
| DREB23_reverse | GCA TCG TTG TTG TTT TGC TCC | |
| FVHsfB3A_forward | GAGCTTACGAGCATGAAAAAC | FVH4_6g24120 |
| FVHsfB3A_reverse | TCTCTTTCTTTCCCTCTCTCC | |
| FvHsfA6a_forward | TCA TCA GCC TAT TCC ACC GC | FvH4_2g23000 |
| FvHsfA6a_reverse | CAT GGC TTG AAG GCA CGC T | |
| FVHsfA1d_forward | GAAGTGCCGAGTGTTAGATG | FVH4_5g22720 |
| FVHsfA1d_reverse | AATGGAGGAAGGAGAAGAGAG | |

**Table S3**. Summary of Differentially methylated regions (DMRs) in strawberry seedling grown at normal and stress conditions

| | | Median | | | Mean | | | | |
|---|---|---|---|---|---|---|---|---|---|
| **Treatment** | **DMR** | **number_C** | **met-diff** | **bps** | **number_C** | **met-diff** | **bps** | **total DMRS** | **Context** |
| Cold | hypermethylated | 12 | 0.14 | 143 | 15 | 0.17 | 182 | 45 | CG |
| Cold | hypomethylated | 12 | -0.13 | 123 | 14 | -0.16 | 157 | 59 | CG |
| Drought | hypermethylated | 10 | 0.17 | 142 | 13 | 0.20 | 162 | 115 | CG |
| Drought | hypomethylated | 12 | -0.23 | 136 | 10 | -0.23 | 165 | 83 | CG |
| Heat | hypermethylated | 10 | 0.17 | 121 | 12 | 0.19 | 140 | 117 | CG |
| Heat | hypomethylated | 12 | -0.15 | 155 | 13 | -0.18 | 183 | 2899 | CG |
| Highlight | hypermethylated | 12 | 0.19 | 136 | 13 | 0.22 | 155 | 81 | CG |
| Highlight | hypomethylated | 12 | -0.25 | 135 | 14 | -0.24 | 161 | 99 | CG |
| Lowlight | hypermethylated | 12 | 0.14 | 129 | 14 | 0.18 | 162 | 110 | CG |
| Lowlight | hypomethylated | 12 | -0.18 | 138 | 14 | -0.20 | 174 | 126 | CG |
| Salicylic | hypermethylated | 12 | 0.14 | 144 | 15 | 0.17 | 163 | 76 | CG |
| Salicylic | hypomethylated | 12 | -0.16 | 122 | 14 | -0.17 | 161 | 109 | CG |
| Salt | hypermethylated | 12 | 0.14 | 131 | 14 | 0.17 | 162 | 74 | CG |
| Salt | hypomethylated | 12 | -0.16 | 143 | 14 | -0.18 | 180 | 357 | CG |
| Cold | hypermethylated | 15 | 0.18 | 279 | 29 | 0.19 | 447 | 8 | CHG |
| Cold | hypomethylated | 26 | -0.20 | 456 | 31 | -0.18 | 482 | 7 | CHG |
| Drought | hypermethylated | 18 | 0.19 | 251 | 22 | 0.20 | 303 | 32 | CHG |
| Drought | hypomethylated | 22 | -0.23 | 243 | 28 | -0.22 | 411 | 23 | CHG |
| Heat | hypermethylated | 19 | 0.21 | 265 | 24 | 0.22 | 335 | 26 | CHG |
| Heat | hypomethylated | 15 | -0.23 | 211 | 19 | -0.23 | 270 | 210 | CHG |
| Highlight | hypermethylated | 21 | 0.23 | 288 | 23 | 0.24 | 342 | 27 | CHG |
| Highlight | hypomethylated | 17 | -0.22 | 256 | 26 | -0.23 | 406 | 26 | CHG |
| Lowlight | hypermethylated | 18 | 0.22 | 179 | 20 | 0.22 | 263 | 23 | CHG |
| Lowlight | hypomethylated | 19 | -0.22 | 212 | 22 | -0.23 | 300 | 38 | CHG |
| Salicylic | hypermethylated | 22 | 0.20 | 282 | 27 | 0.19 | 364 | 28 | CHG |
| Salicylic | hypomethylated | 26 | -0.19 | 303 | 30 | -0.20 | 381 | 16 | CHG |
| Salt | hypermethylated | 24 | 0.17 | 429 | 25 | 0.19 | 388 | 14 | CHG |
| Salt | hypomethylated | 15 | -0.17 | 204 | 20 | -0.19 | 285 | 34 | CHG |
| Cold | hypermethylated | 11 | 0.13 | 40 | 13 | 0.14 | 49 | 1308 | CHH |
| Cold | hypomethylated | 13 | -0.16 | 45 | 15 | -0.16 | 54 | 5041 | CHH |
| Drought | hypermethylated | 13 | 0.14 | 47 | 15 | 0.15 | 58 | 4475 | CHH |
| Drought | hypomethylated | 11 | -0.14 | 41 | 13 | -0.15 | 48 | 1158 | CHH |
| Heat | hypermethylated | 12 | 0.13 | 42 | 14 | 0.14 | 52 | 1037 | CHH |
| Heat | hypomethylated | 13 | -0.17 | 48 | 16 | -0.18 | 58 | 11377 | CHH |
| Highlight | hypermethylated | 13 | 0.16 | 46 | 15 | 0.17 | 57 | 2960 | CHH |
| Highlight | hypomethylated | 12 | -0.14 | 42 | 14 | -0.15 | 51 | 1349 | CHH |
| Lowlight | hypermethylated | 12 | 0.16 | 44 | 14 | 0.17 | 54 | 4775 | CHH |
| Lowlight | hypomethylated | 12 | -0.14 | 42 | 13 | -0.15 | 50 | 2210 | CHH |
| Salicylic | hypermethylated | 12 | 0.15 | 44 | 15 | 0.16 | 53 | 2802 | CHH |
| Salicylic | hypomethylated | 12 | -0.15 | 43 | 13 | -0.16 | 51 | 1266 | CHH |
| Salt | hypermethylated | 12 | 0.13 | 45 | 14 | 0.15 | 53 | 2064 | CHH |
| Salt | hypomethylated | 13 | -0.16 | 45 | 15 | -0.17 | 54 | 4350 | CHH |

**Table S4.** Percentage of genes used for GO enrichment analysis

| Treatment | Genes | Genes with GO number | Genes with no GO number | % Analyzed genes |
|---|---|---|---|---|
| Cold | 2934 | 1545 | 1389 | 52.66 |
| Drought | 2471 | 1267 | 1204 | 51.27 |
| Heat | 6873 | 3765 | 3108 | 54.78 |
| High light | 1804 | 896 | 908 | 49.67 |
| Low light | 2985 | 1553 | 1432 | 52.03 |
| Salt | 3047 | 1614 | 1433 | 52.97 |
| SA | 1820 | 927 | 893 | 50.93 |

**Table S5.** APETALA2/ethylene-responsive element binding protein (AP2/EREBP) superfamily which present DMRs in all different contexts.

| Gene ID | genome | symbol | treatment | meth_diff | DMR size (bp) | context |
|---|---|---|---|---|---|---|
| FvH4_1g16350 | promoter | *AP2-1* | cold | -0.13881 | 40 | CHH |
| | | | heat | -0.162917 | 47 | CHH |
| FvH4_6g34710 | promoter | *AP2-15* | high light | 0.130002 | 63 | CHH |
| FvH4_7g04950 | promoter | *AP2-17* | cold | -0.12849 | 54 | CHH |
| | | | low light | 0.178205 | 32 | CHH |
| FvH4_3g33940 | promoter | *AP2-5* | cold | -0.237724 | 29 | CHH |
| FvH4_2g22290 | promoter | *ERF12* | heat | -0.235965 | 164 | CG |
| FvH4_2g29150 | promoter | *ERF14* | heat | -0.128772 | 70 | CHH |
| | | | salt | -0.147 | 48 | CHH |
| FvH4_2g40810 | promoter | *ERF15* | heat | -0.358406 | 98 | CHH |
| | | | | -0.190833 | 49 | CHH |
| | | | | -0.147292 | 172 | CG |
| FvH4_3g19450 | promoter | *ERF19* | heat | -0.144147 | 23 | CHH |
| FvH4_3g23440 | promoter | *ERF20* | heat | -0.135556 | 85 | CHH |
| | | | | -0.105729 | 96 | CG |
| FvH4_4g03450 | promoter | *ERF22* | heat | -0.245888 | 73 | CHH |
| | | | low light | 0.163993 | 39 | CHH |
| FvH4_4g03470 | gene | *ERF24* | heat | -0.255333 | 115 | CG |
| | | | salt | -0.181481 | 159 | CG |
| FvH4_4g10480 | promoter | *ERF26* | high light | 0.170966 | 69 | CHH |
| FvH4_1g23350 | promoter | *ERF3* | heat | -0.279487 | 73 | CHH |
| | | | low light | 0.206927 | 68 | CHH |
| FvH4_4g22650 | promoter | *ERF30* | heat | -0.416944 | 137 | CG |
| | | | salt | -0.253611 | 137 | CG |
| FvH4_4g27491 | promoter | *ERF32* | heat | -0.227619 | 186 | CG |
| FvH4_5g03840 | promoter | *ERF33* | heat | -0.148553 | 134 | CHH |
| FvH4_5g09520 | promoter | *ERF36* | heat | -0.198205 | 47 | CHH |
| FvH4_2g06050 | promoter | *ERF4* | heat | -0.188491 | 85 | CHH |
| | | | heat | -0.168611 | 79 | CHH |
| | | | Salicylic | 0.138918 | 71 | CHH |
| FvH4_5g19800 | promoter | *ERF40* | salt | -0.187255 | 82 | CHH |
| FvH4_5g19840 | promoter | *ERF41* | heat | -0.209 | 26 | CHH |
| FvH4_6g28880 | promoter | *ERF45* | high light | 0.213818 | 48 | CHH |
| FvH4_6g29930 | promoter | *ERF47* | high light | -0.112958 | 27 | CHH |

| | | | | | | |
|---|---|---|---|---|---|---|
| | | | salt | -0.148333 | 39 | CHH |
| FvH4_6g42000 | promoter | *ERF49* | heat | -0.244 | 49 | CHH |
| | | | heat | -0.122 | 43 | CHH |
| | | | Salicylic | -0.103111 | 43 | CHH |
| FvH4_2g06060 | promoter | *ERF5* | heat | -0.24881 | 57 | CHH |
| | | | heat | -0.12029 | 74 | CHH |
| FvH4_7g10070 | promoter | *ERF53* | salt | 0.145064 | 60 | CHH |
| FvH4_7g15860 | promoter | *ERF55* | heat | -0.175397 | 85 | CHH |
| FvH4_7g26930 | promoter | *ERF59* | heat | -0.266349 | 69 | CHH |
| FvH4_2g13240 | promoter | *ERF6* | heat | -0.279 | 104 | CG |
| FvH4_7g26940 | promoter | *ERF60* | heat | -0.219706 | 43 | CHH |
| | | | salt | 0.175691 | 47 | CHH |
| FvH4_7g30930 | promoter | *ERF61* | salt | 0.147333 | 22 | CHH |
| FvH4_2g21550 | promoter | *ERF7* | heat | -0.29661 | 23 | CHH |
| FvH4_5g01440 | promoter | *FvDREB1* | heat | -0.252986 | 216 | CHH |
| | | | heat | 0.150667 | 39 | CHH |
| | | | salt | -0.205 | 112 | CHH |
| FvH4_5g19440 | promoter | *FvDREB13* | drought | -0.251 | 128 | CG |
| FvH4_6g18090 | promoter | *FvDREB18* | heat | -0.138596 | 68 | CHH |
| FvH4_7g09550 | promoter | *FvDREB20* | heat | -0.134035 | 90 | CHH |
| FvH4_1g09180 | promoter | *FvDREB23* | heat | -0.165238 | 60 | CHH |
| | | | heat | -0.143 | 61 | CHH |
| FvH4_1g16370 | promoter | *FvDREB24* | low light | 0.108581 | 34 | CHH |
| FvH4_5g33180 | promoter | *FvDREB27* | high light | 0.209042 | 36 | CHH |
| FvH4_1g21210 | promoter | *FvDREB29* | heat | -0.336667 | 118 | CG |
| | | | | -0.262745 | 90 | CHH |
| FvH4_5g37820 | promoter | *FvDREB31* | heat | -0.370333 | 83 | CG |
| | | | | -0.17534 | 88 | CG |
| | | | | -0.120926 | 130 | CHH |
| FvH4_6g26090 | promoter | *FvDREB32* | drought | 0.237534 | 27 | CHH |
| | | | heat | -0.178805 | 28 | CHH |
| | | | heat | -0.178739 | 33 | CHH |
| FvH4_6g43870 | promoter | *FvDREB7* | cold | -0.292202 | 28 | CHH |
| | | | heat | -0.186389 | 21 | CHH |
| | | | heat | -0.179399 | 30 | CHH |
| FvH4_3g44200 | promoter | *RAV1* | heat | -0.190302 | 104 | CHH |
| | | | salt | -0.149703 | 69 | CHH |
| FvH4_6g45390 | promoter | *RAV7* | heat | -0.150833 | 76 | CHH |
| | | | heat | -0.142 | 51 | CG |

| | | | | | | |
|---|---|---|---|---|---|---|
| | | | high light | -0.19079 | 43 | CHH |

**Table S6.** Heat shock transcription factors (HSFs) which contains DMRs under stress.

| Gene ID | genome | symbol | treatment | meth_diff | DMR size (bp) | context |
|---|---|---|---|---|---|---|
| FvH4_5g01770 | promoter | FvHsfB2b | heat | -0.2475 | 116 | CG |
| | promoter | | heat | -0.230725 | 97 | CHH |
| FvH4_6g22550 | - | FvHsfA4b | - | - | - | - |
| FvH4_4g13230 | promoter | FvHsfA5a | heat | -0.112075 | 132 | CG |
| FvH4_3g07170 | promoter | FvHsfA4a | cold | -0.184167 | 43 | CHH |
| FvH4_2g23000 | promoter | FvHsfA6a | heat | -0.191714 | 186 | CG |
| FvH4_4g01180 | promoter | FvHsfA6a | heat | -0.178667 | 42 | CHH |
| FvH4_4g33360 | promoter | FvHsfA8a | heat | -0.168614 | 57 | CHH |
| FvH4_3g09340 | - | FvHsfB2a | - | - | - | - |
| FvH4_7g29190 | TES | FvHsfB2a | heat | -0.183001 | 258 | CHG |
| | TES | | heat | -0.158826 | 32 | CHH |
| FvH4_2g34260 | - | FvHsfB4a | - | - | - | - |
| FvH4_1g04810 | promoter | FvHsfC1a | heat | -0.16278 | 46 | CHH |
| FvH4_5g22720 | promoter | FvHsfA1d | cold | -0.105129 | 44 | CHH |
| | promoter | | heat | -0.109583 | 66 | CHH |
| | promoter | | heat | -0.105714 | 43 | CHH |
| FvH4_1g07800 | promoter | FvHsfA9a | drought | 0.123981 | 133 | CHH |
| | promoter | | SA | 0.23 | 37 | CHH |
| FvH4_2g31690 | - | FvHsfA2a | - | - | - | - |
| FvH4_6g17890 | - | FvHsfA3a | - | - | - | - |
| FvH4_6g24120 | promoter | FvHsfB3a | heat | -0.123865 | 160 | CG |
| FvH4_3g16090 | promoter | FvHsfA1b | cold | -0.147773 | 167 | CHH |
| | promoter | | heat | -0.300605 | 41 | CHH |
| | promoter | | salt | -0.13751 | 154 | CHH |
| FvH4_1g16030 | promoter | FvHsfB1a | salt | 0.127333 | 67 | CHH |
| | promoter | | salt | 0.122637 | 38 | CHH |
| FvH4_6g27900 | promoter | FvHsfA4c | cold | -0.212994 | 37 | CHH |
| | promoter | | cold | -0.231754 | 34 | CHH |
| | TES | | heat | -0.153333 | 124 | CG |
| | promoter | | drought | 0.326217 | 167 | CHH |
| | promoter | | low light | 0.272328 | 48 | CHH |

| | | | | |
|---|---|---|---|---|
| promoter | salt | 0.107894 | 82 | CHH |

**Table S7.** Association of stress-indeced differentially methylated regions with transposable elements in F. vesca

| TE | TOTAL counts in genome |
|---|---|
| DNA/DTA | 10867 |
| DNA/DTC | 14437 |
| DNA/DTH | 6119 |
| DNA/DTM | 27122 |
| DNA/DTT | 1190 |
| DNA/Helitron | 19743 |
| long_terminal_repeat | 2434 |
| LTR/Copia | 12750 |
| LTR/Gypsy | 12690 |
| LTR/unknown | 27460 |
| MITE/DTA | 2293 |
| MITE/DTC | 225 |
| MITE/DTH | 2289 |
| MITE/DTM | 5898 |
| MITE/DTT | 7 |

(a) Total number of TE in the *Fragaria vesca* genome

| TE family | cold | drought | heat | high_light | low_light | SA | salt |
|---|---|---|---|---|---|---|---|
| DNA/DTA | 512 | 470 | 974 | 314 | 546 | 347 | 504 |
| DNA/DTC | 500 | 463 | 998 | 429 | 649 | 373 | 510 |
| DNA/DTH | 253 | 242 | 650 | 166 | 299 | 160 | 251 |
| DNA/DTM | 1215 | 1158 | 2366 | 804 | 1343 | 807 | 1203 |
| DNA/DTT | 45 | 34 | 121 | 33 | 43 | 22 | 41 |
| DNA/Helitron | 504 | 427 | 1294 | 374 | 533 | 309 | 516 |
| LTR/Copia | 421 | 373 | 817 | 307 | 440 | 312 | 445 |
| LTR/Gypsy | 546 | 549 | 1021 | 446 | 690 | 422 | 611 |
| LTR/unknown | 1313 | 1323 | 2464 | 1007 | 1535 | 987 | 1536 |
| MITE/DTA | 143 | 95 | 292 | 92 | 151 | 79 | 142 |
| MITE/DTC | 12 | 5 | 24 | 3 | 6 | 1 | 7 |
| MITE/DTH | 154 | 100 | 327 | 95 | 148 | 65 | 133 |
| MITE/DTM | 391 | 347 | 727 | 206 | 413 | 222 | 370 |
| MITE | 700 | 547 | 1370 | 396 | 718 | 367 | 652 |

(b) Number of TEs with differentially methylated regions.
